# Electrostatic Tuning of a Potassium Channel in Electric Fish

**DOI:** 10.1101/206243

**Authors:** Immani Swapna, Alfredo Ghezzi, Michael R. Markham, D. Brent Halling, Ying Lu, Jason R. Gallant, Harold H. Zakon

## Abstract

Molecular and biophysical variation contributes to the evolution of adaptive phenotypes, particularly behavior, though it is often difficult to understand *precisely* how. The adaptively significant electric organ discharge behavior of weakly electric fish is the direct result of biophysical membrane properties set by ion channels. Here we describe a voltage-gated potassium channel gene in African mormyrid electric fishes, that is under positive selection and highly expressed in the electric organ. The channel produced by this gene shortens electric organ action potentials by activating quickly and at hyperpolarized membrane potentials. Surprisingly, the source of these unique properties is a derived patch of negatively charged amino acids in an extracellular loop near the voltage sensor. Further, we demonstrate that this portion of the channel functions differently in vertebrates than the generally accepted model based on the *shaker* channel, and suggest a role for this loop in the evolutionary tuning of voltage-dependent channels.

## Introduction

A major goal of evolutionary biology is understanding how changes at the genetic level result in adaptations. Much can be learned about the genetic basis of adaptation by studying species with unique and/or extreme phenotypic adaptations (1–5). Because electric fish signal with electricity, and those signals are under strong selection and generated by ion channels, electric fish are an excellent system to study how selection may tune ion channels and, especially, how it might push them towards their biophysical limits.

Nocturnally active mormyrid electric fish produce brief, weak voltage electric fields, called electric organ discharges (EODs) to form electrosensory images of their environments and to communicate during courtship (6). Though EOD waveforms are known to be influenced by both natural and sexual selection pressures (7–9), the molecular targets and biophysical mechanisms for selection remain elusive. The superfamily Mormyroidea is composed of the Gymnarchidae and the Mormyridae (Fig. 1). The family Gymnarchidae is composed of a single species, *Gymnarchus niloticus* that generates a constant sine wave-like EOD. The sister taxon, the Mormyridae (blue triangle, Fig. 1), includes hundreds of species all of which produce brief, often multi-phasic pulses at variable intervals--“pulse-type” EODs (10).

**Figure 1.**
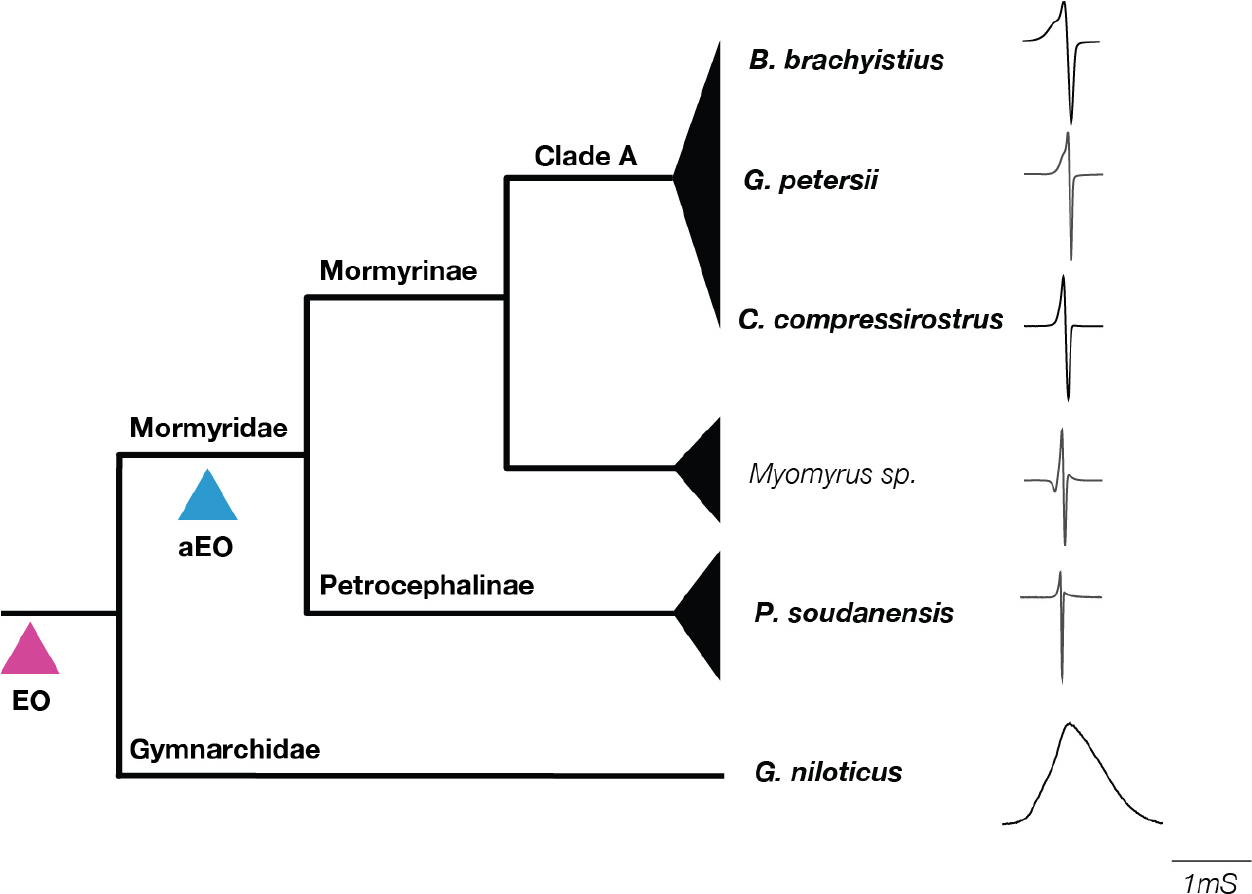
Major clades of mormyroidea with representative electric organ discharges (EODs). Species used in this study are shown in boldface. Electric organs (EO; purple triangle) evolved in the common ancestor of the superfamily mormyroidea that is comprised of two families, the monotypic “Gymnarchidae” and the “Mormyridae.” Electric organ discharges in *Gymnarchus niloticus* are quasi-sinusoidal wave-like discharges, although only a single cycle is shown here. In the common ancestor of the Mormyridae, a more specialized adult electric organ (aEO, blue triangle) evolved, which produces short-duration pulse type discharges. EODs of Petrocephalinae (e.g. *Petrocephalus soudanensis*) and non-clade A mormyrinae (e.g. *Myomyrus* spp.) are characteristically short duration, whereas clade-A mormyrinae are highly diverse in EOD properties, including duration and number of phases (complexity). Some EOD recordings shown derive from specimens deposited in the Cornell Museum of Vertebrates and EOD recordings deposited in the Macaulay Library at the Cornell Laboratory of Ornithology: *Myomyrus macrops* (CUMV 92394; ML EOD 515280), *Brienomyrus brachyistius* CUMV 80464; ML EOD 510794) *Gnathonemus petersii* (CUMV 87880; ML EOD 510874). EOD from *Petrocephalus soudanensis* was provided by C.D. Hopkins (Cornell University), and EODs from *Gymnarchus niloticus* and *Campylomormyrus compressirostrus* were recorded by J. Gallant from captive laboratory specimens.

In the Mormyridae, selection has often favored the evolution of extremely brief EODs (11, 12) and neuronal adaptations to discriminate between timing differences less than 10 microseconds (13, 14), suggesting EOD pulse duration must be precisely regulated. EODs are produced by the synchronous discharge of action potentials (APs) from many hundreds of individual electrocytes. These APs are extremely brief (~200 microseconds) (15, 16). Because the duration of action potentials is shaped by potassium currents, we assessed whether voltage-gated potassium channels expressed in the mormyrid electric organ (EO) have evolved specializations for generating brief APs. Here we report a potassium channel gene with a strong imprint of positive selection that is expressed at high levels in the EO. The biophysical properties of the channel encoded by this gene are specialized for generating brief action potentials. These properties result from an evolutionarily novel motif of negatively charged residues in an extracellular loop adjacent to the channel’s voltage-sensor. The previous understanding of this extracellular loop is largely based on the *shaker* potassium channel in fruit flies. However, our analyses suggest that this loop functions differently in vertebrates and suggests a role for this loop in the evolutionary tuning of ion channels.

## Results

### Identification of a rapidly evolving, electric organ-expressing potassium channel

We began by examining muscle and EO transcriptomes representative of the Gymnarchidae and the Mormyridae (Tables S1,2). We uncovered two orthologs of the mammalian *KCNA7* channel gene--*kcna7a* and *kcna7b*--that duplicated in the teleost whole genome duplication (Fig. 2A, S1). In mammals, KCNA7 is expressed in heart and muscle (17–19); *kcna7b* is also expressed in muscle in mormyrids (Fig. 2B). While *kcna7a* is expressed in muscle in *Gymnarchus*, it is predominantly expressed in the EO in mormyrids (Fig. 2B) where it is the major potassium channel gene expressed, and without a beta subunit (Fig. 2C). The change in expression from muscle to EO in the ancestral mormyrid was accompanied by a burst of positive selection (HYPHY, REL branch-site model (20)) (p<0.01, corrected p value) on this branch (Fig. 2A, 2D).

**Figure 2.**
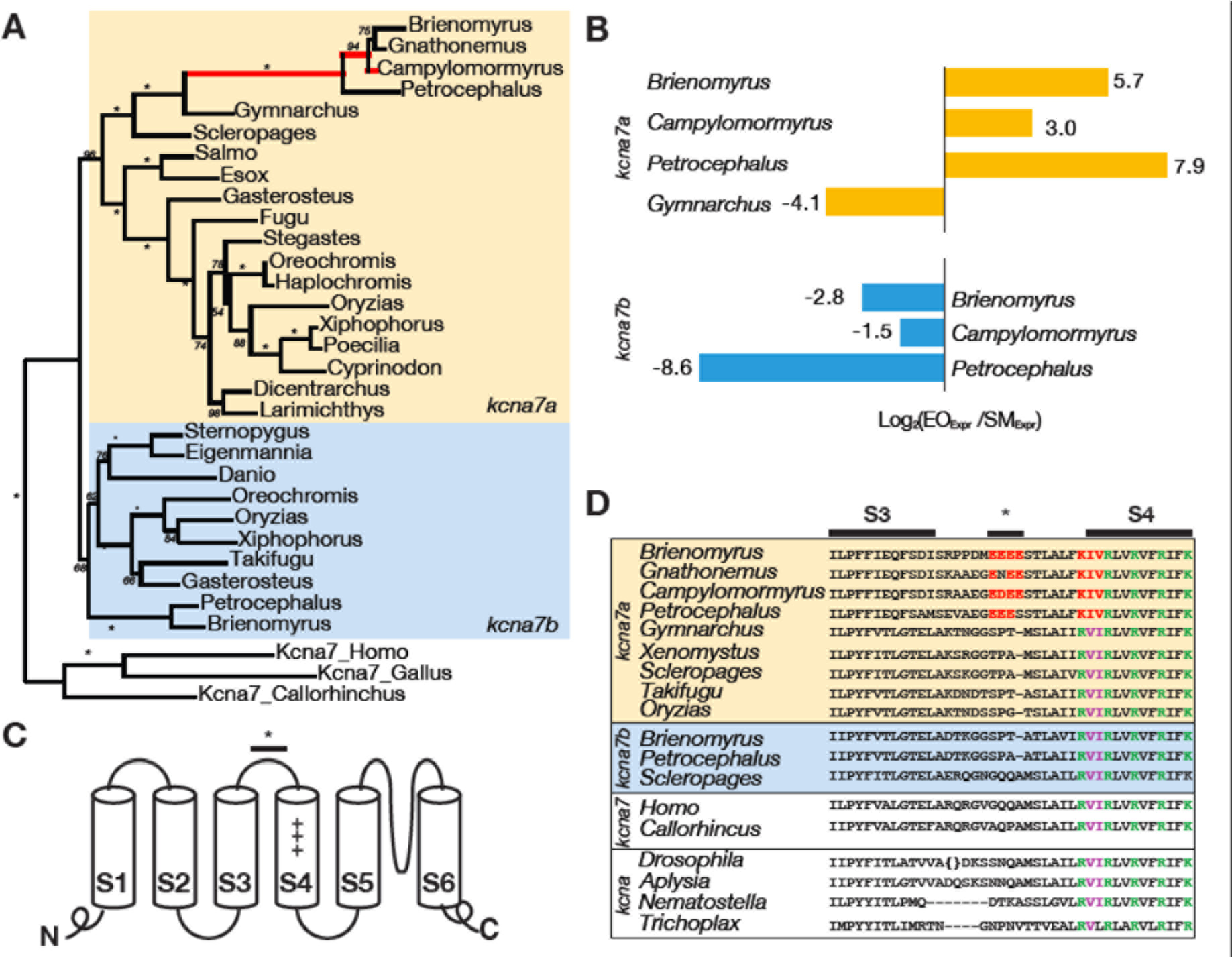
Evolutionary change in electric organ-expressing potassium channels of mormyrid electric fish. (A) Maximum likelihood gene tree for *kcna7a* and *kcna7b* illustrates the duplication of *KCNA7* in teleosts (asterisks = bootstrap values of 100). Branches displaying episodic diversifying selection are in red (p<0.01, corrected p-value). *Petrocephalus* is a basal group and the other three species are in the derived “clade A” giving good phylogenetic coverage of the mormyrids. Note the burst of episodic selection at the base of pulse-generating mormyrids and within “clade A.” (B) Relative expression of *kcna7a* and *kcna7b* in the skeletal muscle (SM) and electric organ (EO) of mormyrids and *Gymnarchus*. (C) Schematic illustration of potassium channel. Note the voltage sensor (S4) and the S3-S4 linker. A bar represents the portion of the S3-S4 linker that is the focus of our study. The asterisk over the bar indicates the location of a site determined to be under positive selection by HYPHY. The amino acid sequences of this region are shown in (D). (D) S3-S4 linker and S4 in *shaker* channels: *kcna7a* of four mormyrids (*Brienomyrus*, *Gnathonemus*, *Campylomormyrus*, *Petrocephalus*), *Gymnarchus*, and other teleosts (*Xenomystus*, *Scleropages*, *Fugu*, *Oryzias*); kcna7b of mormyrids and another teleost, and *KCNA7* in human (*Homo*) and elephant shark (*Callorhinchus*); the single *shaker* channel in fruit fly (*Drosophila*) and sea slug (*Aplysia*), and one of six *shaker* channels in sea anemone (*Nematostella*). Conserved positively charged amino acids in S4 in blue, conserved hydrophobic amino acids in S4 in violet. Amino acid substitutions in S4 and the S3-S4 linker in mormyrids in red. The S3-S4 linker of the *Drosophila* shaker gene is longer than in vertebrates (see: Fig. S6) and is truncated to fit the alignment (parentheses). Complete species names and accession numbers for all sequences are given in table S5.

### Voltage-clamp analysis and modeling show biophysical specializations for generating brief action potentials

A potassium current can shorten action potentials if it activates quickly or at a relatively hyperpolarized membrane potential, or has a steep conductance-voltage curve. In *Xenopus* oocytes, we expressed both *kcna7a* sequence from *Gymnarchus niloticus,* which expresses kcna7a in muscle and *kcna7a* sequence from *Brienomyrus brachyistius*, a mormyrid which expresses *kcna7a* in EO and has a brief EOD pulse. We observed that the potassium current from *Brienomyrus* activates faster (tau: *Brienomyrus* = 1.2 +/− 0.16 msec; *Gymnarchus* = 3.04 +/− 1.0 msec, p<0.0001, Fig. 3A-C), at ~30 mV more hyperpolarized membrane potentials (V_1/2_: *Brienomyrus* = −44.4 +/− 5.7 mV; *Gymnarchus* −13.1+/− 5.9 mV, p<0.0001, Fig. 3D,E), and with a steeper conductance-voltage (G-V) curve (k: *Brienomyrus* = 7.0 +/− 1.4; *Gymnarchus* 9.3 +/− 2.3; p <0.028, Fig. 3F) than the current from *Gymnarchus*.

**Figure 3.**
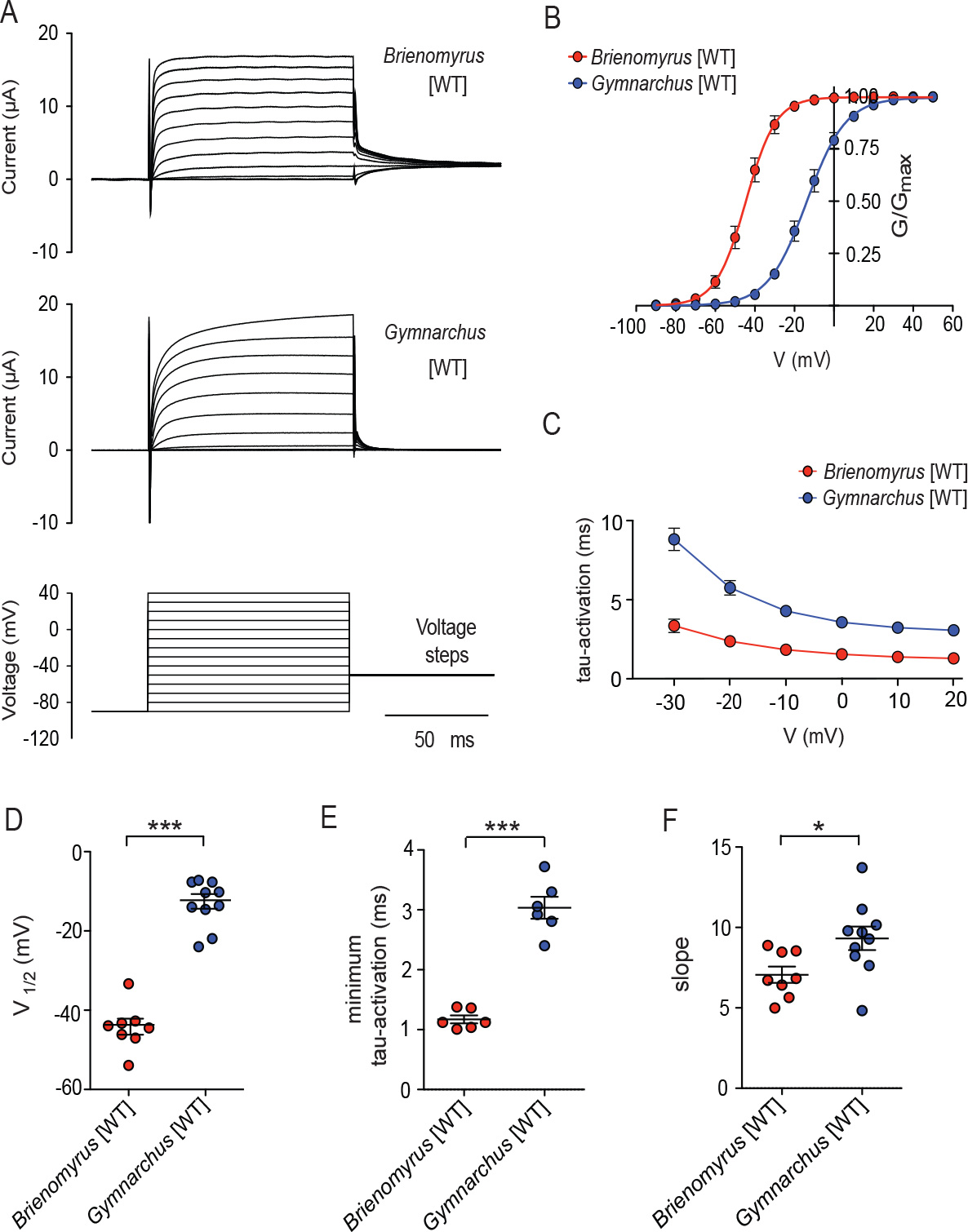
Currents from *kcna7a* of *Gymnarchus* and *Brienomyrus* expressed in *Xenopus* oocytes show striking differences. (A) Raw voltage-clamp traces (B) conductance-voltage (G-V) curves (C) time constant of activation (tau) as a function of voltage (D) the voltage at which the conductance is 50% of maximum (V_1/2_) (E) minimum tau (taken at +20 mV) (F) the slope of the G-V curve. Note that the expressed current from *Brienomyrus* activates faster, at more hyperpolarized voltages, and with a steeper slope that that from *Gymnarchus*. Here and in all subsequent figures, *Gymnarchus* = blue; *Brienomyrus* = red. * = p<0.05, *** = p<0.001 (unpaired *t* test).

Next, we constructed a computational model of electrocyte action potentials, using the parameters from these recordings, together with generic sodium currents (Fig. 4). We found that changes in the G-V curve slope did not significantly affect AP width, but that the tau-V curve, G-V curve midpoint (V_1/2_), and Na^+^ current duration all contributed in changes in AP width (R^2^ = 0.94, df = 4, 14576, p < 0.001). Within this regression model the tau-V curve accounted for ~29% of AP width variance, G-V curve midpoint accounted for ~25% of AP width variance, and Na^+^ current duration accounted for ~40% of AP width variance. Taken together, these results strongly suggest that differences in K^+^ conductance properties between *G. niloticus* and *B. brachyistus* exert a major influence on AP width.

**Figure 4.**
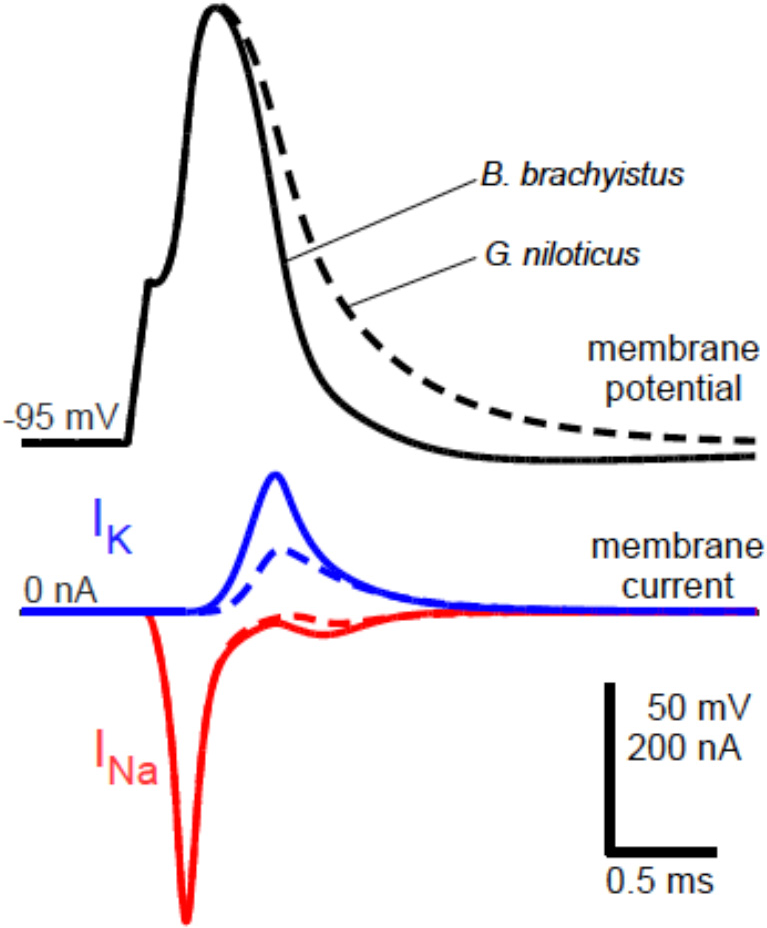
Computational simulations of electrocyte action potentials. Model cells included a generic Na^+^ conductance, linear leak, and a voltage-gated K^+^ conductance. The K^+^ conductance was modeled using parameters derived from experimental data for either *Brienomyrus* Kv1.7a or *Gymnarchus* Kv1.7a. Stimulus current was a 0.1-ms 150 nA step pulse. The model cell with *Brienomyrus* Kv1.7a exhibits shorter action potential duration than the model cell with *Gymnarchus* Kv1.7a.

### Site-directed mutagenesis identifies a novel functional motif

Potassium channels have six transmembrane helices, S1-S6, in the alpha subunit (Figure 2C). Four alpha subunits combine to form a channel. S5 and S6 line a conductive pore for potassium that opens when membrane depolarization alters the conformations of S4, which forms the voltage sensor. The *kcna7a* gene, which encodes the Kv1.7a potassium channel protein, is a member of the *shaker* family of potassium channels named after the canonical *shaker* gene of *Drosophila*. The *shaker* family is ancient preceding the divergence of bilateria and cnidaria (21). We constructed alignments of Kv1.7 channel proteins with their orthologs including distantly related *shaker* family channels (Fig 2D). These protein alignments highlight that, despite strong conservation of the amino acids in the S4 over ~800 million years (22), three amino acid substitutions occurred in the S4 of the ancestor of the pulse mormyrids following the mormyrids’ divergence from *Gymnarchus* and before their radiation (23) (Fig. 2D).

We tested whether these amino acid substitutions alter channel properties by a swapping amino acid residues reciprocally between the *Gymnarchus* and *Brienomyrus* Kv1.7a proteins using site-directed mutagenesis. Placing the three amino acids from the S4 of *Brienomyrus* (Fig. 2D) into the S4 of *Gymnarchus* produced a modest shift in the expected direction (leftward) of the G-V curve, and complex shifts in activation kinetics suggesting that amino acids at other sites interact with these to influence tau-activation (Fig. 5). The complementary substitutions from *Gymnarchus* to *Brienomyrus* had no effect on voltage sensitivity and only minor effects on tau-activation (Fig. 6). From these results, we concluded that other amino acid substitutions accrued during the evolution of the ancestral pulse mormyrid Kv1.7a, and the effects of these substitutions must override the effects of the substitutions in S4.

**Figure 5.**
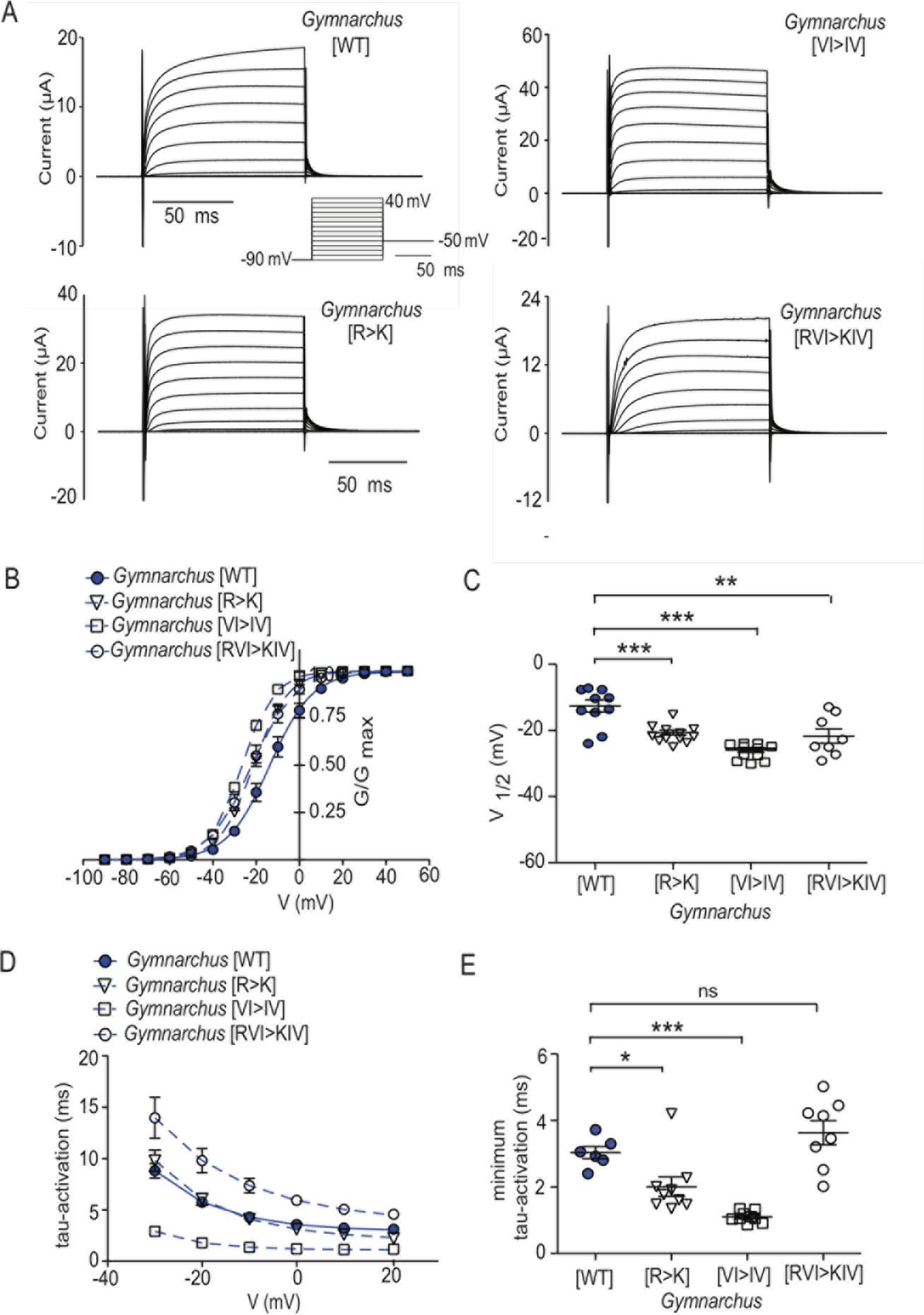
Replacement of first three amino acids of *Brienomyrus* S4 into *Gymnarchus* S4 (R>K; VI>IV; RVI>KIV). (A) Raw data (B) G-V curves (C) V_1/2_ (D) tau-activation (E) minimum tau-activation.

**Figure 6.**
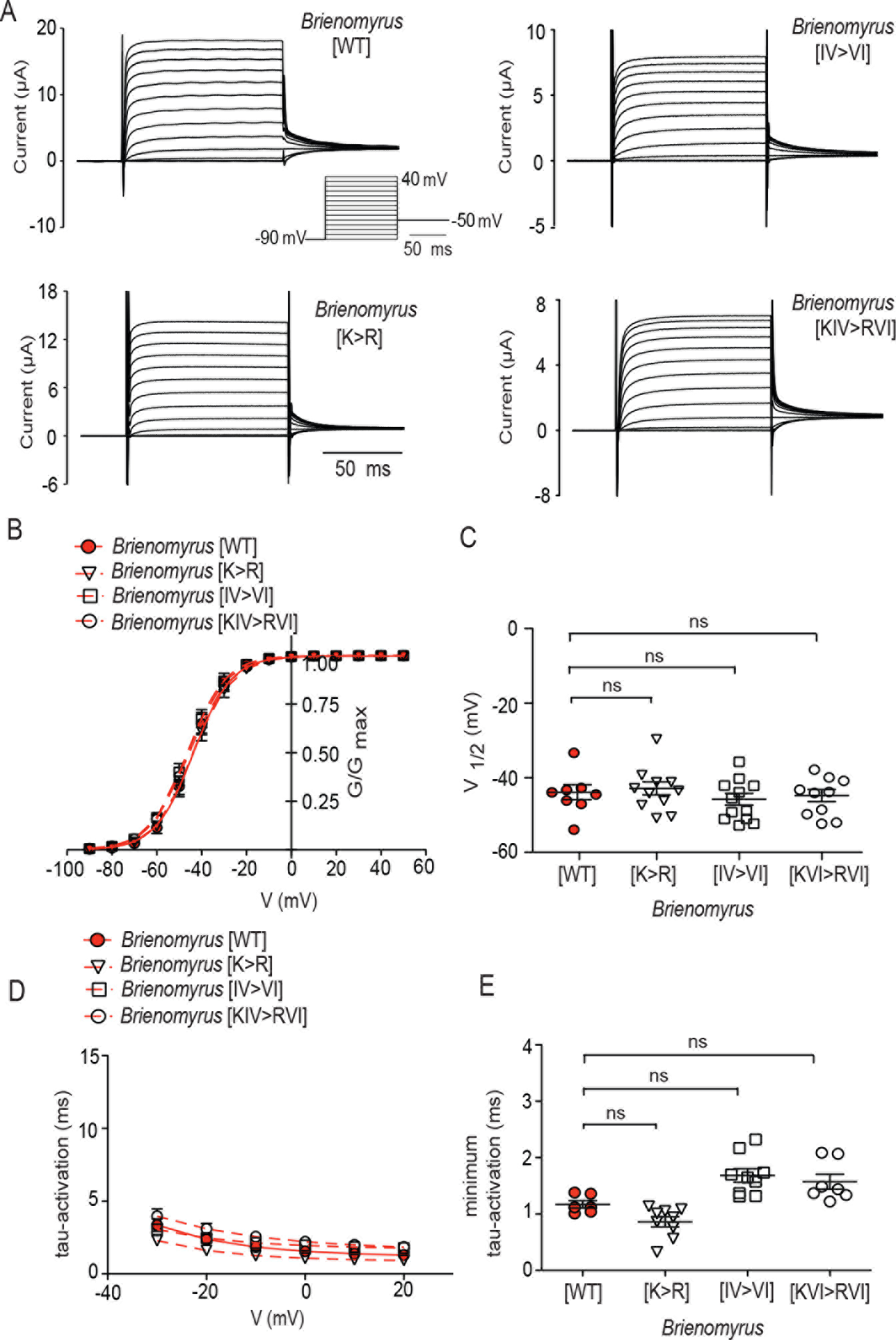
Replacement of first three amino acids of *Gymnarchus* S4 into *Brienomyrus* S4 (K>R; IV>VI; KIV>RVI). (A) Raw data (B) G-V curves (C) V_1/2_ (D) tau-activation (E) minimum tau-activation.

The mormyrid kv1.7a is unusual among vertebrate voltage-gated potassium channels at another place: it possesses a patch of contiguous negatively charged amino acids in the S3-S4 linker (Fig. 2D). A site within this patch was detected by HYPHY as evolving under positive selection (asterisk, Fig. 2C,D). Again, using site-directed mutagenesis, we swapped homologous regions of the S3-S4 linker between *Gymnarchus* and *Brienomyrus* channels. This resulted in a strong shift in the V_1/2_ and tau-activation of one species to that of the other (*Brienomyrus* [EEEE>SPT] = −9.34 +/− 4.6 mV; *Gymnarchus* [SPT>EEEE] = −40.69 +/− 4.7 mV) (WT vs. chimeric channel in both species, p<0.0001) (Fig. 7).

**Figure 7.**
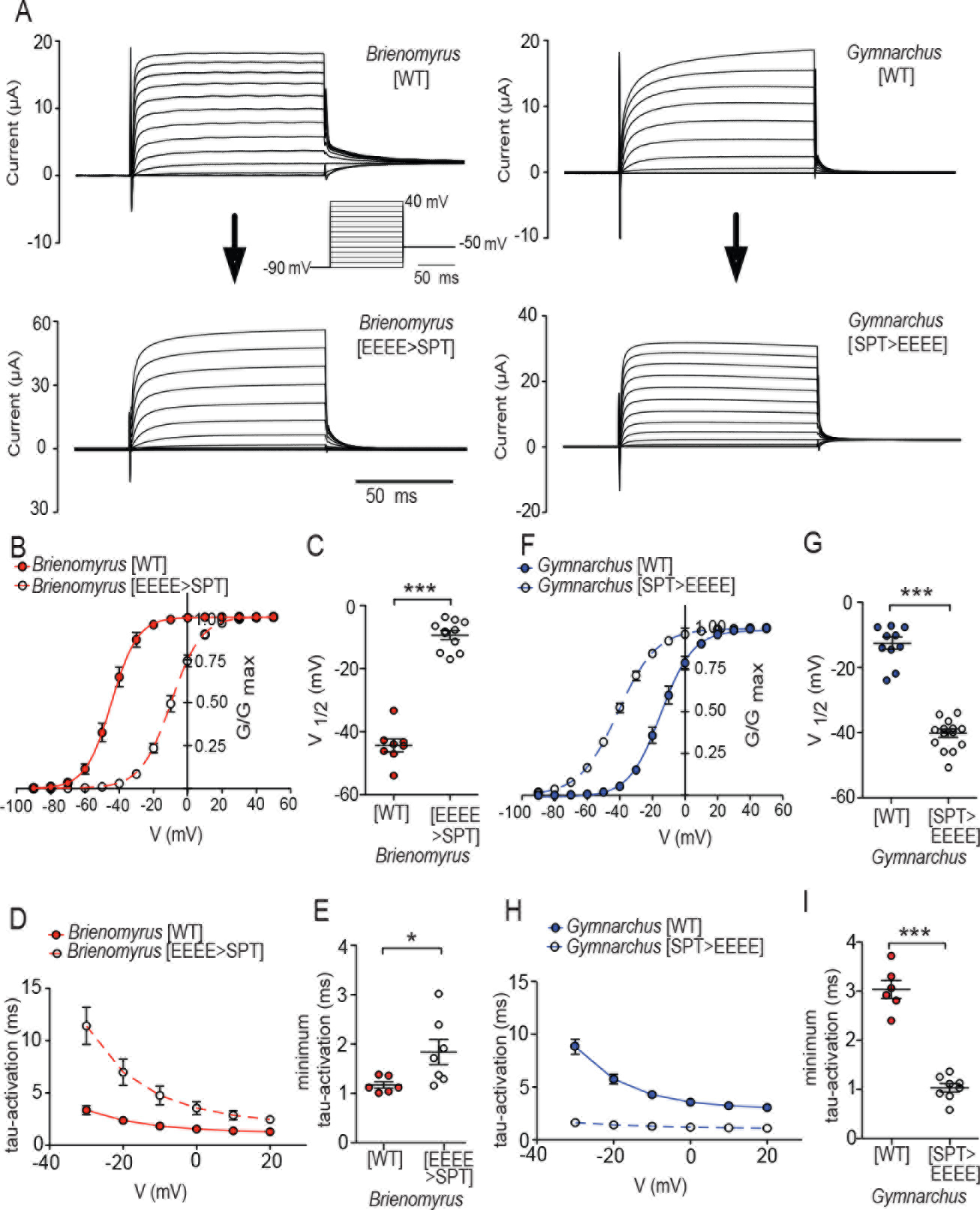
The distinctive characteristics of the currents from *kcna7a* of *Gymnarchus* and *Brienomyrus* are transferred by swapping part of the S3-S4 linker. (A-E) currents of *Brienomyrus* wild type (WT) and *Brienomyrus* (EEEE>SPT); (A) raw currents (B) G-V curves (C) V_1/2_ (D) tau-activation (E) minimum tau-activation; (F-J) currents of *Gymnarchus* WT and *Gymnarchus* (SPT>EEEE) (F) G-V curves (G) V_1/2_ (H) tau-activation (I) minimum tau-activation. Here and in all subsequent figures, solid lines = WT; dotted lines = chimeric channels. * p<0.05, *** p<0.001 (unpaired *t* test).

### Predicted structural properties of S3-S4 linkers

There is an increasing recognition that the S3-S4 linker influences potassium channel properties (24). The prevailing view is that the S3-S4 linker’s influence on *Drosophila shaker* channel properties is due to its length and flexibility rather than its charged residues (25–27). However, we note the S3-S4 linker of the *shaker* channel of *Drosophila* and other arthropods is atypically long compared with that of other animals (Fig. S2), suggesting that S3-S4 linker may function differently in arthropods than in other animals. Therefore, we made a structural model of the Kv1.7a S3-S4 linkers of *Brienomyrus* and *Gymnarchus*.

The extra negative charges in the S3-S4 linker of *Brienomyrus* could contribute to different physical properties between the channels of the two species, including flexibility and charge. Although no structures are available for Kcna7a (Kv1.7), voltage sensors from other channels have been determined (28–32). Models were generated based on homology to structures of these voltage sensors in the Protein Data Bank in order to qualitatively compare the differences between the *Gymnarchus* and the *Brienomyrus* Kcna7a voltage sensors (Fig. 8). Consistent with a flexible S3-S4 linker, different modelers returned varying solutions for its structure, whereas portions of the transmembrane helices were consistently modelled (Fig. 8A,B). The lengths of TM3 and TM4 are not well determined, and different models had different solutions for the extent of helical content of the S3-S4 linker.

Although these generated models may not represent actual physiological states, we can map the physical properties that amino acid sequence confers to general positions in structure space (33). To compare how the physical properties of amino acids can differentiate the two voltage sensors, we used the voltage sensor predictions that were most alike between *Brienomyrus* and *Gymnarchus*. First, we compared flexibility. Amino acids that are mostly found in flexible protein domains are more concentrated in the S3-S4 linker in the models of both voltage sensors (Figs. 8C,D). Residues that may add rigidity to the loops are found at different locations (Fig. 8G). Taken as a whole, the overall content of flexible voltage-sensor residues is comparable.

Finally, the presence of four negatively charged glutamates is expected to contribute to a strong surface charge on the protein in the S3-S4 linker of *Brienomyrus* compared to that of *Gymnarchus* (Fig. 8E,F). The strong negative charge on *Brienomyrus* is nearly sufficient to cap the voltage sensor, whereas the negative surface charge on *Gymnarchus* covers a much smaller area. Compared to flexibility, the differences in protein surface charge appear to be much greater at distinguishing the differences between these voltage sensors. The models emphasize an electrostatic influence of the negative charges in the *Brienomyrus* S3-S4 linker on channel gating.

### Electrostatic tuning of the voltage sensor

To test whether the additional glutamates act *via* their electrostatic properties, we conducted two experiments: replacing negatively charged aspartates for glutamates did not alter V_1/2_ (V_1/2_ [EEEE>DDDD] = −49.2 +/− 4.9 mV) and replacement with positively charged amino acids produced an even more profound rightward shift of voltage sensitivity than replacement with *Gymnarchus*’ neutral amino acids (V_1/2_ [EEEE>KKKK] = +16.0 +/−9.2 mV) (Fig. 9).

**Figure 8.**
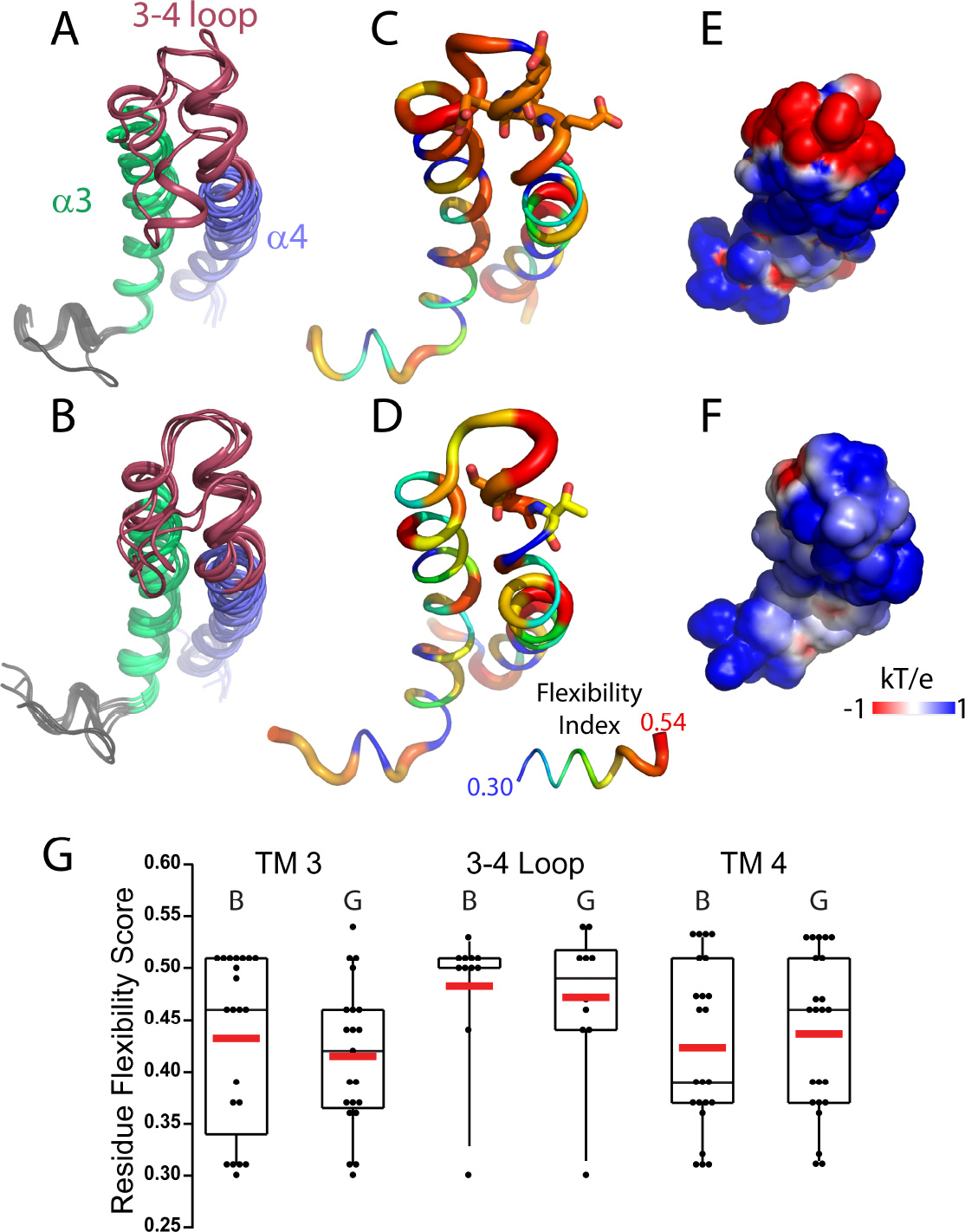
Comparison of voltage sensor of Kcna7a from *Brienomyrus* (A, C, and E) and *Gymnarchus* (B, D, and F). Panels A and B show the best solutions for several different structure prediction algorithms. Each best solution is depicted as a cartoon ribbon that traces the voltage sensor backbone. Transmembrane helices are colored green and blue for α3 (S3) and α4 (S4) respectively. The S3-S4 loop is colored burgundy. In C and D, the flexibility score of a residue was mapped to a putty scheme of the structural models that are most alike between *Brienomyrus* and *Gymnarchus*. The inset under D shows that thicker, redder putty depicts amino acids that are more frequently found in flexible protein domains, whereas thinner and bluer are mostly found in rigid domains. Comparing the same models as in C and D, Panels E and F show the electrostatic surface charge of the voltage sensors, with the range of charge shown in the inset in the lower right of F. (G) Standard box plots of voltage sensor domain flexibility by residue of *Brienomyrus* B; and *Gymnarchus*, G. Data are included as black dots representing amino acid flexibility scores for residues in transmembrane helix 3 (TM 3), the 3-4 loop, and TM4. The mean of each plot is included as a red bar.

**Figure 9.**
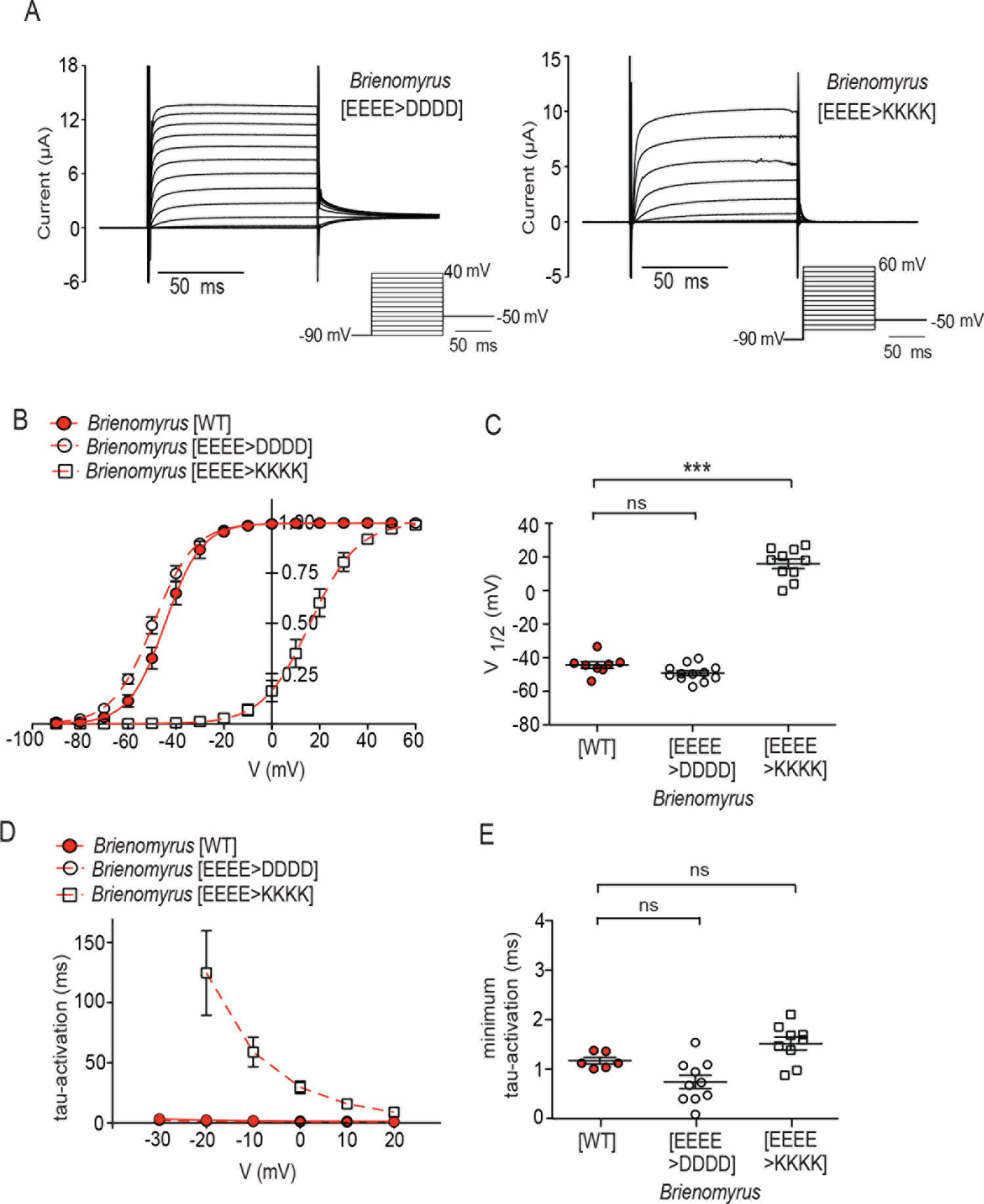
Substitution of negatively charged (D) but not positively charged (K) amino acids in the *Brienomyrus* S3-S4 linker retains WT biophysical properties. (A) Representative currents (B) conductance-voltage curves (C) V_1/2_ values (D) tau-activation curves (E) minimum tau-activation curves. ns p> 0.05, *** p<0.001 (one-way ANOVA followed by Dunnett’s post-test compared to (34)).

How might the negative patch influence channel behavior? The “repulsion hypothesis” suggests that this patch might face another negatively charged amino acid or patch of amino acids in the closed state. Under these conditions a focused, local interaction between the two could create a repulsive bias so that less depolarizing voltage is needed to move the S4 (35). Alternatively, the negatively charged amino acids in this region may be forming salt bridges with positively charged amino acids in other parts of the channel favoring the open confirmation. The “surface charge” hypothesis states that the charged intra- and extracellular parts of proteins globally add to, or cancel out, the membrane potential caused by separation of ions. In this way S4 movement could occur at less depolarized membrane potentials.

To distinguish between these hypotheses, we devised a test based on the premise that if a local bias occurs *via* a local electrostatic repulsion, it should not matter if interacting partners are negative or positive. We identified negatively charged putative interaction partners near the S3-S4 linker (S3-S4, S1-S2, S5-pore), converted them to positive residues (E/D>K), and tested whether this altered V_1/2_ (Figs. S3-5). Some substitutions had little effect but some, such as D379K, made V_1/2_ more positive. We then combined any substitutions that shifted V_1/2_ with the positively charged [EEEE>KKKK] S3-S4 linker. We observed that the combination of two positive patches only shifted V_1/2_ to even more positive values, never a reversion to a more negative V_1/2_. Substituting the positive amino acids in the extracellular loop regions to negative amino acids did not have any effect on the V_1/2_. These results suggest that the negative patch in the S3-S4 linker is not acting *via* a local repulsive force or formation of salt bridges but, rather, globally by contributing to the surface charge.

## Discussion

The *kcna7a* potassium channel gene evolved rapidly preceding the adaptive radiation of the mormyrids. This is a similar pattern to that of a muscle-expressing sodium channel gene that also shifted its expression to the mormyrid EO (*scn4aa*) (36, 37). The resulting amino acid substitutions in the S3-S4 linker in Kv1.7a contribute to biophysical changes that enable ultra-brief EODs. While there are a number of other amino acid substitutions at conserved sites, future studies will elucidate their functions. We note that a few species of mormyrids, such as *Campylomormyrus tshokwe,* have secondarily and independently evolved long-duration EOD pulses (8, 38, 39). The expression level of some Kv1 family genes (40) have been implicated in species differences in EOD duration in the explosively radiating genus *Campylomormyrus*, though *kcna7a* paralogs have not been examined in this group. It will be interesting to examine whether additional novel amino acid substitutions have occurred in *C. tshokwe kcna7a*, and within other species with long duration EOD pulses.

The prevailing view of the S3-S4 linker’s influence on *shaker* channel properties is that it is due to its length and flexibility rather than its charged residues (25–27). Our data show that the long S3-S4 linker of the *Drosophila shaker* and other arthropods is not representative of other animal groups, including vertebrates (Fig. S2). The S3-S4 linker of mormyrids possesses a naturally-occurring, extreme but instructive case of a large patch of negatively charged amino acids poised directly above the S4; other vertebrate potassium channels have negatively charged amino acids at one or more sites within the S3-S4 linker that could mediate such effects. Our study and a few other recent studies (35, 41) suggest that negative charges acting to alter outer membrane surface charge rather than flexibility are at play in tuning the voltage-sensitivity of vertebrate voltage-gated potassium channels. This concept is further supported by a complementary observation of an L-type Ca^2+^ channel found in elasmobranch electroreceptors that activates at more hyperpolarized potentials than expected due to a patch of positively charged amino acids in the S2-S3 linker localized near the inner mouth of the channel (42). Thus, it seems that the addition of charged amino acids on either side of the S4 voltage sensor may be an evolutionary simple and widespread way to modify channel voltage-sensitivity and gating kinetics.

## Acknowledgements

The sequence data reported here are available from NCBI (accession numbers given in supplementary materials). Funding for this work was by NSF IOS # 1557657 (JRG), NSF IOS# 1557857 (HHZ), NSF IOS# 1350753 and IOS1257580 (MRM), NIH 2R01NS077821, to Richard Aldrich).

The authors declare no competing interests.

## Materials and Methods

### Animal Sources

*<u>Petrocephalus soudenensis</u>* [Osteoglossiformes: Mormyridae]– A freshwater mormyrid species native to central Africa was obtained through the aquarium trade. Sex of individual was not determined. Cornell Museum of Vertebrates (CUMV: 91327, Specimen # 5727; Identified by M.E. Arnegard). Tissues were collected in RNA later prior to RNA-isolation.

*<u>Gymnarchus niloticus</u>* [Osteoglossiformes: Gymnarchidae]) – A freshwater species native to central Africa, was obtained through the aquarium trade. Sex of individual was not determined. Tissues were collected in RNA later prior to RNA-isolation.

All procedures used followed the American Physiological Society Animal Care Guidelines, and were approved by the Institutional Animal Care and Use Committee at University of Texas, Austin and Michigan State University.

### RNA Extraction and Library Preparation for RNAseq

Various mRNA libraries were sequenced on various platforms as summarized in Supplemental Table S1.

*<u>RNA Sequencing</u>* – Tissues were homogenized in liquid nitrogen using a ceramic mortar and pestle, and total RNA was extracted using Trizol (ThermoFisher Scientific, Waltham, MA USA) following the manufacturer’s specifications (see (43) for detailed methods). Total RNA was quantified using qubit and quality assessed using a Bioanalzyer (Agilent Technologies, Santa Clara, CA USA). Samples were then depleted of ribosomal RNA using the a RiboZero kit (Illumina, Inc. San Diego, CA) as per manufacturer’s specifications, or RNA samples were again assessed for concentration and quality before preparation of sequencing libraries. cDNA libraries were constructed from ribosomal RNA-depleted samples using the Illumina TruSeq RNA Sample Preparation (v.2) kit. All libraries were sequenced on an Illumina HiSeq2500 using 125bp single-end reads (2x125bp).

### Additional Data Sources

*<u>B. brachyistius</u>* – We downloaded the raw reads for *Brienomyrus brachyistius* electric organ and skeletal muscle tissues referenced in (1) with NCBI BioProject accession # (PRJNA248545).

*<u>C. compressirostrus</u>* – We downloaded the raw reads for skeletal muscle and electric organ tissue referenced in (44) with NCBI BioProject accession # (PRJNA192446).

### Transcriptome Assembly

For each species, short read sequences obtained from each tissue were combined.
Quality control, adapter trimming, and quality filtering was performed using Trimmomatic version 0.33 (45) with the following settings: ILLUMINACLIP TruSeq3-PE.fa:2:30:10 SLIDINGWINDOW:4:30 MINLEN:60. Paired trimmed reads for each species were then assembled into de-novo transcriptomes using Trinity v.20140413p1 (46) with default parameters. To speed up assembly, reads from *C. compressirostrus* were digitally normalized prior to assembly using the --normalize_reads flag. Results of all assemblies are summarized in Table S2.

### Calculation of Expression Levels

Short reads from individual tissue libraries were mapped to the appropriate transcriptome assembly using Bowtie2 v. 2.2.6 (47) with default parameters. Read counting and ambiguity resolution were performed using RSEM v. 1.3.0, (47). Raw read counts were then scaled and normalized per sample by transforming to FPKM values, and samples were cross-sample normalized using a trimmed mean of M values (TMM; (48)) for comparison between libraries and species. Scaling and normalization was performed using Trinity v.20140413p1 scripts ‘`align_and_estimate_abundance.pl`’ `and` ‘`abundance_estimates_to_matrix.pl`’ `respectively`.

### Assignment of Orthologues Between Electric Fish

Using SeaView (http://doua.prabi.fr/software/seaview) nucleotide or amino acid sequences of mormyrid potassium channel genes were aligned with potassium channel genes (in the *shaker* family: nucleotide = kcnax; amino acid = kv1.x) of other teleosts, and human. Trees were rooted with *D. melanogaster shaker*, the canonical *shaker* family channel. Alignments were analyzed in a maximum likelihood format.

### Data Availability

The raw sequence data generated in this project are available through the NCBI Sequence Read Archive (SRA) with the following accession numbers. Assembled transcriptomes and expression tables are available for download and BLAST searching via the EFISHGENOMICS web-portal (http://efishgenomics.integrativebiology.msu.edu). Focal sequences described in this analysis have been additionally deposited in NCBI Genbank with the accession numbers described in table S3.

### Mutagenesis and Expression in Oocytes

The coding sequences of Kcna7a genes from *Brienomyrus brachyistius* and *Gymnarchus niloticus* were synthesized by GenScript (Piscataway, NJ, USA) and cloned into the pGEMHE vector. During synthesis the Kozak sequence (GCCACC) was added to the 5’ end of the sequences immediately before the start codon to improve translation efficiency.

Mutations were introduced using the QuikChange II XL site directed mutagenesis kit from Agilent technologies, USA, and verified by sequencing. *In-vitro* transcription was carried out using the mMESSAGE mMACHINE T7 transcription kit from Thermo Fisher Scientific as per the manufacturer’s protocol. The concentration of the synthesized capped mRNA was measured using the ND-1000 spectrophotometer from NanoDrop Technologies, Wilmington, DE, USA. Each oocyte was injected with 50 nl of 0.01-0.1 μg/μl mRNA solution. Injected oocytes were incubated at 16°C in sterile modified Barth’s solution [88 mM NaCl, 1 mM KCl, 2.4 mM NaHCO3, 19 mM HEPES, 0.82 mM MgSO4, 0.33 mM Ca(NO3)2, 0.91 mM CaCl2, 10,000 units/l penicillin, 50 mg/l gentamicin, 90 mg/l theophylline, and 220 mg/l sodium pyruvate, pH 7.5] for 1-3 days after injection to achieve optimal channel expression for recordings.

### Electrophysiology and Analysis

Currents from mRNA injected oocytes was recorded using the two-electrode voltage clamp. All recordings were done at room temperature (21-23 C) using the OocyteClamp OC 725C amplifier from Warner Instruments Corp (Hamden, CT, USA). The bath solution contained 115 mM NaCl, 1.5 mM KCl, 10 mM HEPES and 1 mM MgCl_2_, pH- 7.4 adjusted with NaOH.The pipette solution consisted of 3 M KOAc and 15 mM KCl. Current activation was measured using a series of 10mV voltage steps (100 ms each) from a potential of −90mV to 40mV followed by a 100 ms tail pulse of −50 mV. For recording currents from *Brienomyrus* mutants that open at more depolarized potentials the voltage steps used were −70 to 60 mV or 80 mV in 10 mV increments. The K^+^ reversal potential was measured by a pulse protocol of 100 ms depolarization to 40 or 60 mV followed by a 200 ms test pulse of −120 mV to 0 mV in 10 mV increments. In all recording the holding potential was maintained at −90 mV.

Data acquisition and the preliminary minimal analysis of I-V data were done using the pCLAMP 8 software from Axon Instruments, Inc., (Foster City, CA, USA). All other analysis was done using the GraphPad Prism 5 software. The ionic current (I) recorded during the voltage steps was converted to conductance (G) by dividing with the driving force (V-E_rev_). The half-activation potential (V_1/2_) and slope factor of the activation curve were obtained by fitting the G-V curves with a simple Boltzmann function. For calculation of tau-activation of the rising phase of the K^+^ channel current at each voltage step was fitted with the equation I*(t*)= Imax*(1−exp(−K*t))^n where K= 1/tau-activation. Although, we tried fitting with n=2,3,4, the best overall fit was obtained with n=2 and this has been used throughout this study.

Statistical analysis was done using the GraphPad Prism 5 software. Statistically significance was determined using the student’s t test or one-way ANOVA followed by the Dunnett’s post-test comparing to the WT with a 95% confidence interval.

### Computational Methods

For numerical simulations we modeled the electrocyte as a single compartment. The capacitance *C* was 30 nF consistent with empirical measurements of whole-cell capacitance in electrocytes of weakly electric fish (49–51). Differential equations were coded and integrated with Matlab (Mathworks, Inc., Natick MA) using Euler’s method with integration time steps of 1 × 10^−9^ sec. The current balance equation was:

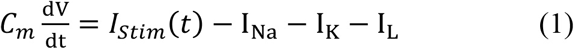

where I_Na_ represents Na^+^ current, I_K_ represents a non-inactivating delayed rectifier K^+^ current, and I_L_ is the leak current. Equations for these currents were as follows:

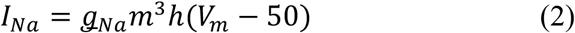

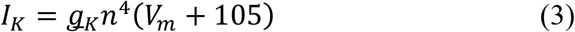

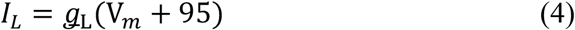

The gating variables in Equations 2 and 3 are given by Equation 5 where j = m, h, or n:

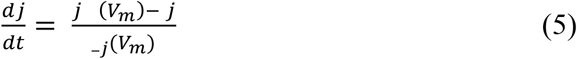

The voltage-dependent values of *j* evolved as follows:

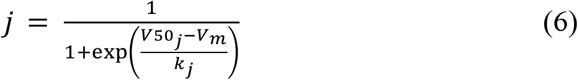

where *V*50_*j*_ and *k*_*j*_ are derived from Boltzmann sigmoidal fits to empirical from the present results for j = n and from previous empirical data for electrocyte Na^+^ conductances for j = m (51).

These values are given in Table 1. t_*j*_ is given by Eqn. 7 for j = m and n:

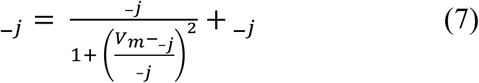

and by Eqn. 8 for j = h:

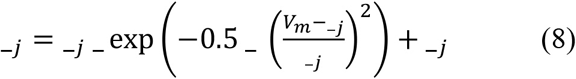

Where values of α_*j*_, β_*j*_, μ_*j*_, and σ_*j*_, were determined by least-squares best fits to empirical data from the present results for j = n, and from previous empirical data (52) for j = m or h. These parameter values are given in Table S1.

We first compared simulated Aps from model cells with identical Na^+^ and leak conductance parameters, but K^+^ conductance parameters derived from G-V and τ-V curves for kcna7a of either *G. niloticus* or *B. brachyistus* (Supplemental Figure 2). To further explore the contributions of these parameters to changes in AP width, we tested a set of 14,580 model cells where we systematically varied k_n_, V50_n_, and μ_n_ between the values for *G. niloticus* and those for *B. brachyistus*, thereby changing the G-V curve slope, G-V curve midpoint, and τ-V curve, respectively. We also assessed the potential effect of increasing Na^+^ current duration by increasing _–h_ from 0.08 ms to 0.16 ms in steps of 0.01 ms, thereby slowing Na^+^ inactivation (parameter ranges are shown in Table S2). We then used stepwise multiple linear regression (Matlab *stepwiselm* function) to determine the relative contributions of these four parameters in determining AP width across all combinations of these parameters.

### Structure Models

The Protein Modeling Portal, http://www.proteinmodelportal.org (53) was used to submit projects to multiple servers for structure prediction. Kcna7a voltage-sensor domain protein sequences were submitted using *Brienomyrus* (residues 276-340, accession #####) and *Gymnarchus* (residues 275-337, accession #####). Some solutions returned obvious steric clashes. As a final step, the Yasara server was used to energy minimize solutions, add hydrogens, and to alleviate steric clashes within the models (http://www.yasara.org/minimizationserver.htm)(54).

The values used for amino acid flexibility score were determined previously (55), and rendered in place of B-factors in the model output structure files. Electrostatic surface potential was determined using the PDB2PQR and APBS (Adaptive Poisson-Boltzmann Solver) plugins for The PyMOL Molecular Graphics System, Version 0.99rc6 (Schrödinger, LLC) (56, 57).

## Supplementary Figures

**Figure S1.**
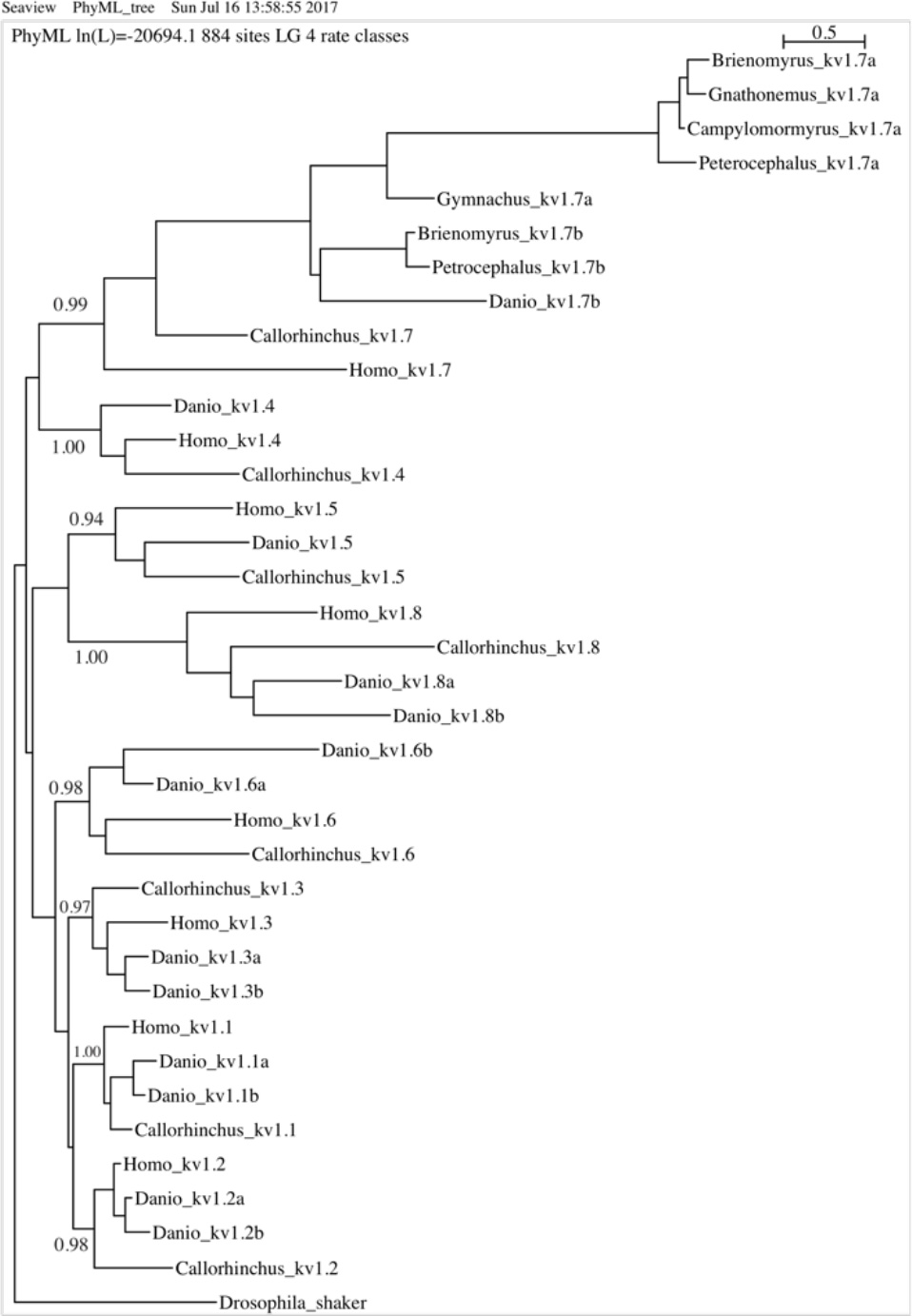
Phylogenetic tree of vertebrate Kv1 (*shaker*) family channels (bootstrap support given for the node for each channel, nomenclature based on mammalian kv1 channels, rooted with *Drosophila melanogaster shaker*). Note strong bootstrap support for the placement of mormyroid kv1.7a and kv1.7b paralogs with those of other vertebrates. Mormyridae = *Brienomyrus brachyistius*, *Campylomormyrus compressirostris*, *Gnathonemus petersii*, *Petrocephalus soudanensis*. Gymnarchidae = *Gymnarchus niloticus*. Other teleostei = *Danio rerio*. Chondrichthyes = *Callorhinchus milli*. Mammalia = *Homo sapiens*. Accession numbers in Table S5.

**Figure S2.**
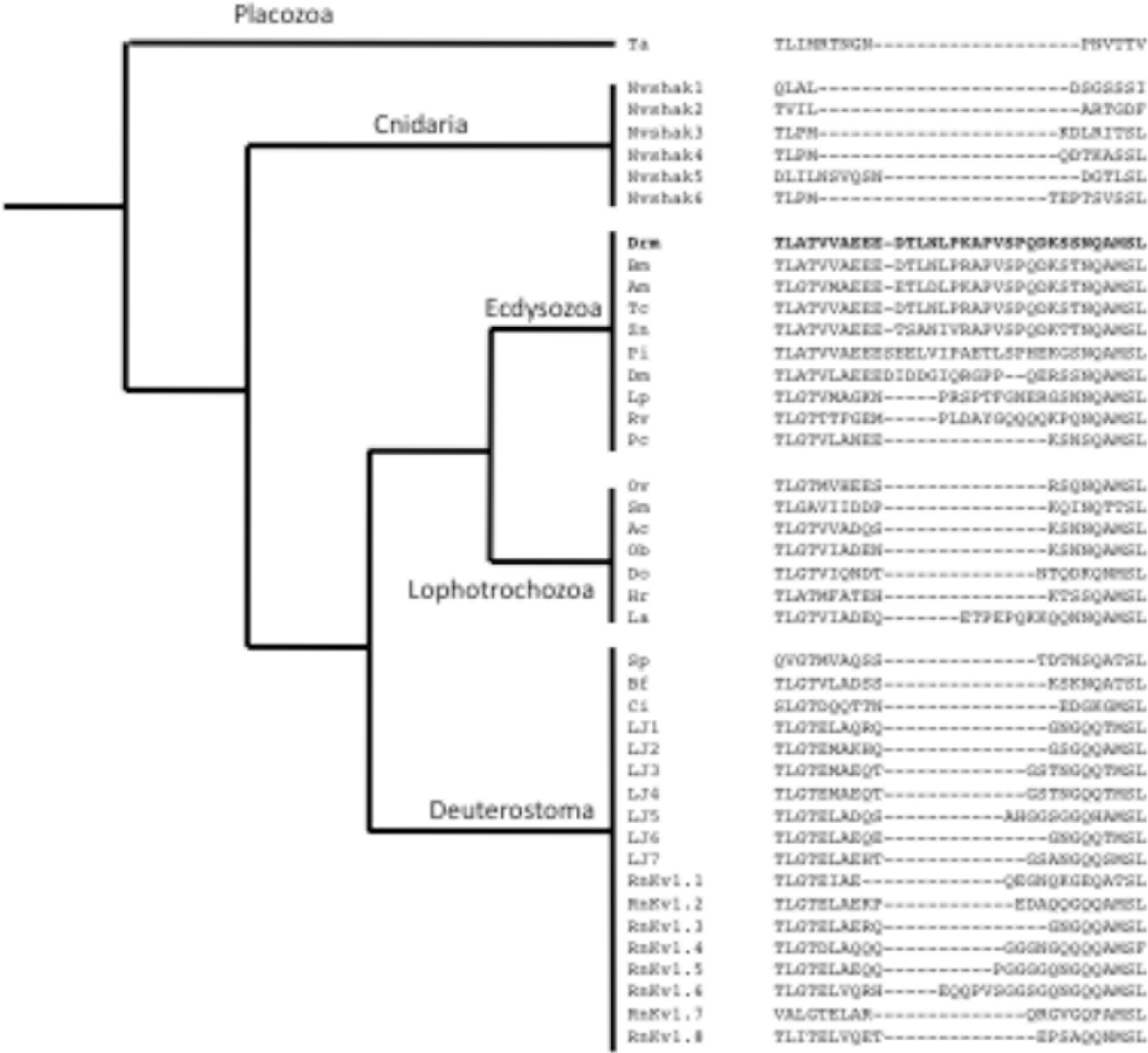
Schematic phylogeny of the S3-S4 linker in the *shaker* family of potassium channels of animals rooted with placozoan shaker. Note that the S3-S4 linker is long only in arthropods. The canonical *shaker* channel of *Drosophila melanogaster* is **bold**. Ta = the placazoan, *Trichoplax adherens*; Nv = starlet sea anemone, *Nematostella vectensis* (six shaker paralogs, shak1-6); Drm = fruit fly *Drosophila melanogaster*; Bm = silk moth, *Bombyx mori*; Am = honey bee, *Apis mellifera*; TC = red flour beetle, *Tribolium castaneum*; Zn = dampwood terminte, *Zootermopsis nevadensis*; Pi = spiny lobster, *Panulirus interruptus*; Dm = water flea, *Daphnia magna*; Lp = horseshoe crab, *Limulus polyphemus*; Nc = golden orb-web spider, *Nephila clavata*; Rv = water bear, *Ramazzottius varieornatus*; Pc = penis worm, *Priapulus caudatus*; Ov = liver fluke, *Opisthorchis viverrini*; Sm = trematode, *Schistosoma mansoni*; Ac = California sea hare, *Aplysia californica*; Ob = California two spot octopus, *Octopus bimaculoides*; Do = opalescent inshore squid, *Doryteuthis opalescens*; Hr = leech, *Helobdella robusta*; La = brachiopod, *Lingula anatine*; Sp = purple sea urchin, *Strongylocentrotus purpuratus*; Bf = the Florida lancelet or *Amphioxus, Branchiostoma floridae*; Ci = sea squirt, *Ciona intestinalis*; Lj = Japanese lamprey, *Lethenteron japonicum* (seven *shaker* paralogs); Rn = brown rat, *Rattus norvegicus* (eight *shaker* paralogs (kv1.1-kv1.8).

**Figure S3.**
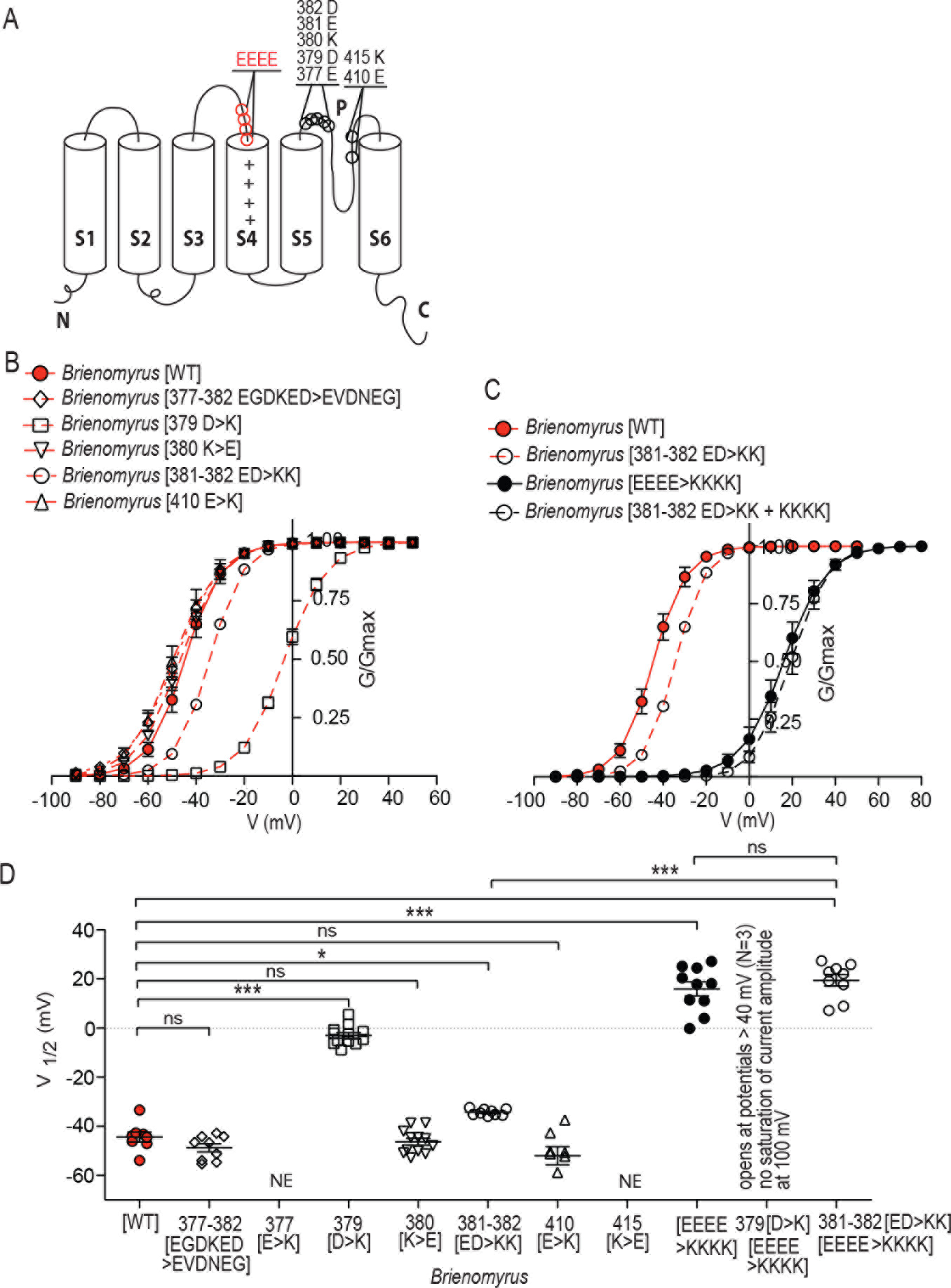
Conversion of charged amino acids to opposite charge in S5-pore region alone and in combination with S3-S4 EEEE>KKKK in *Brienomyrus* kv1.7a. (A) Schematic diagram of site of amino acid substitutions in S5-pore (black) and S3-S4 (red) (B) G-V curves for amino acid substitutions alone (C) G-V curves for amino acid substitutions in S5-pore that affected V_1/2_ in combination with S3-S4 substitutions. (D) Summary of V_1/2_ values for all experimental groups. NE = not expressing.

**Figure S4.**
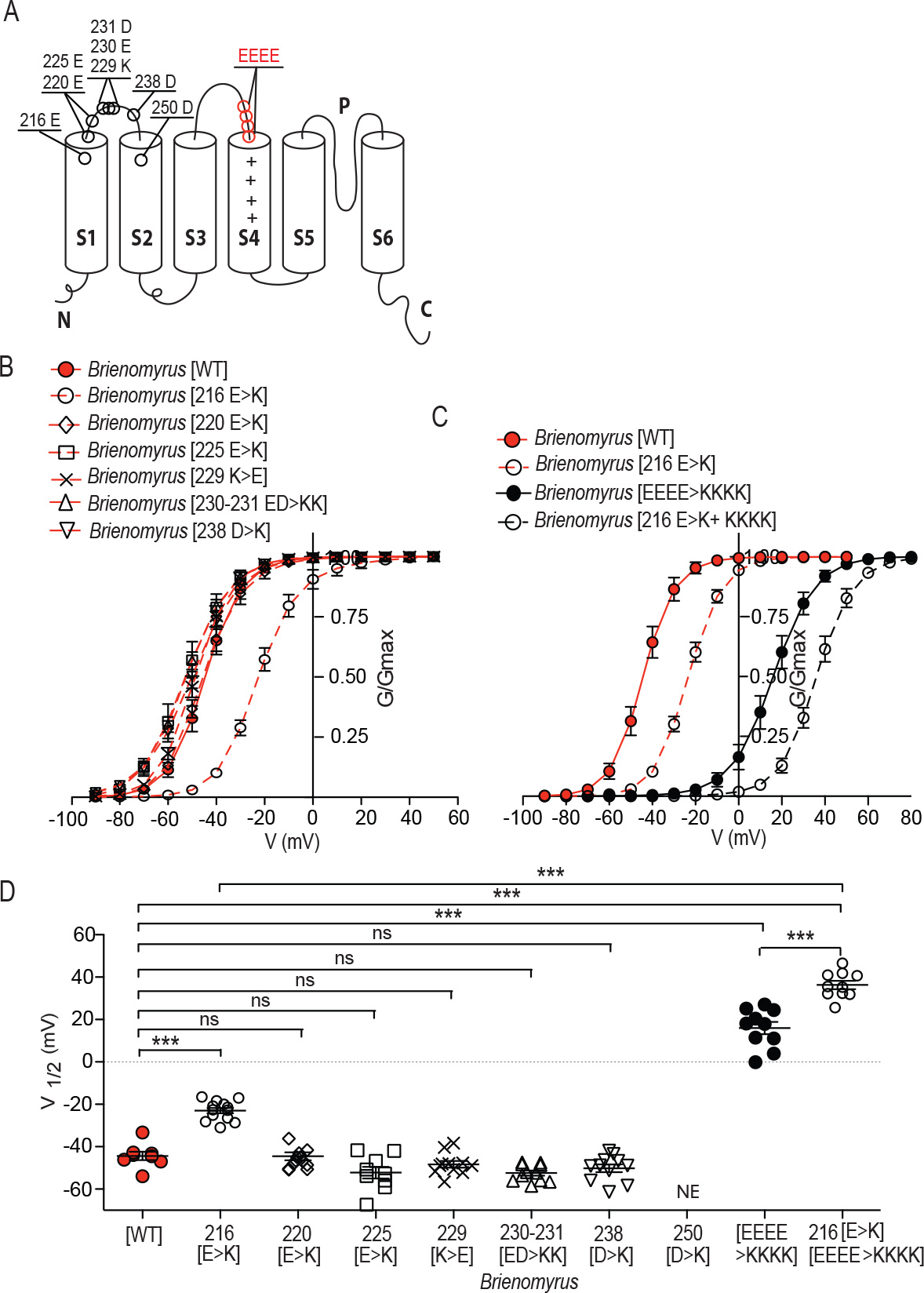
Conversion of charged amino acids to opposite charge in S1-S2 loop alone and in combination with S3-S4 EEEE>KKKK in *Brienomyrus* kv1.7a. (A) Schematic diagram of site of amino acid substitutions in S1-S2 loop (black) and S3-S4 (red) (B) G-V curves for amino acid substitutions alone (C) G-V curves for amino acid substitutions in S1-S2 loop that affected V_1/2_ in combination with S3-S4 substitutions. (D) Summary of V_1/2_ values for all experimental groups. NE = not expressing.

**Figure S5.**
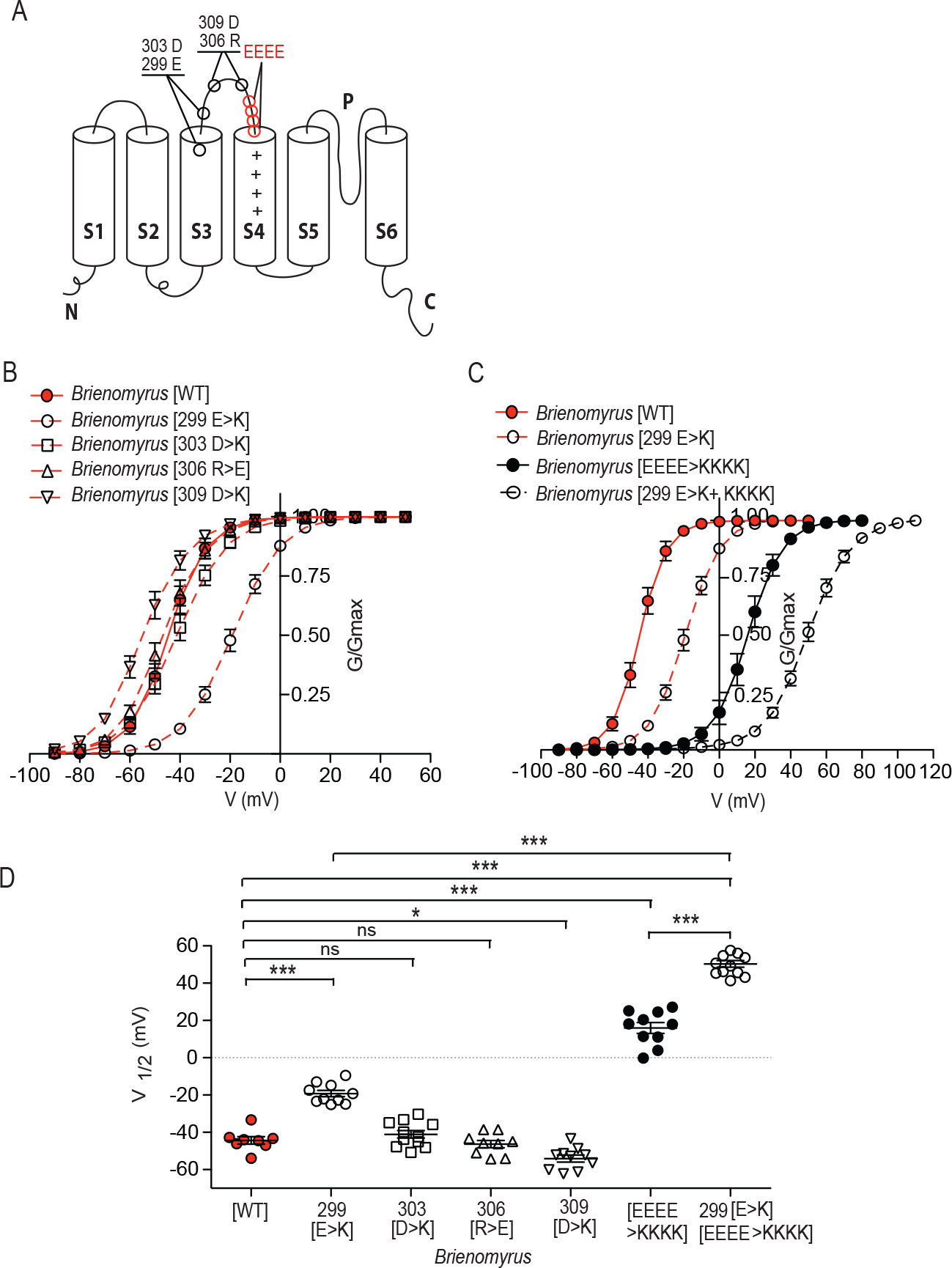
Conversion of charged amino acids to opposite charge in S3 and proximal S3-S4 loop alone and in combination with S3-S4 EEEE>KKKK in *Brienomyrus* kv1.7a. (A) Schematic diagram of site of amino acid substitutions in proximal S3 and proximal S3-S4 loop (black) and S3-S4 (red) (B) G-V curves for amino acid substitutions alone (C) G-V curves for amino acid substitutions in S3 and proximal S3-S4 loop that affected V_1/2_ in combination with S3-S4 substitutions. (D) Summary of V_1/2_ values for all experimental groups.

**Table S1.** Supplemental Table 1 - Summary of RNA-Seq Data Collected.

| Species | NCBI BioProject | NCBI SRA | samples | sequencing |
| --- | --- | --- | --- | --- |
| *Petrocephalus soudenesis* |  |  | 3 tissues – brain, electric organ, skeletal muscle | Illumina Hi |
| *Gymnarchus niloticus* |  |  | 2 tissues – electric organ, skeletal muscle | Illumina Hi |

**Table S2.**
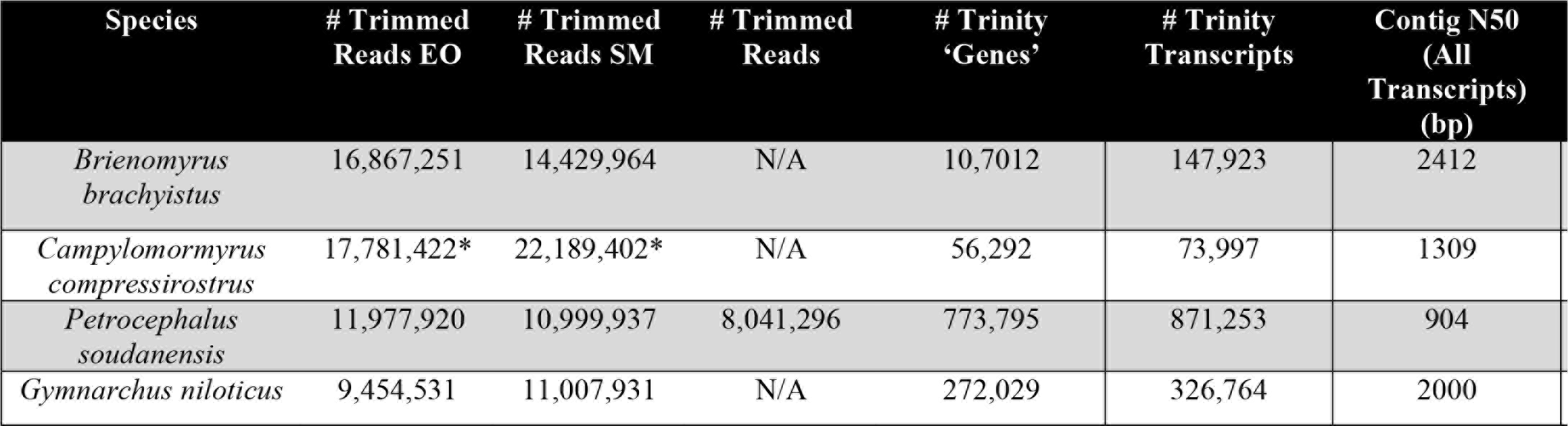
Supplemental Table 2 - Summary Statistics of Trimmed Reads and Transcriptome Assemblies.

**Table S3.**
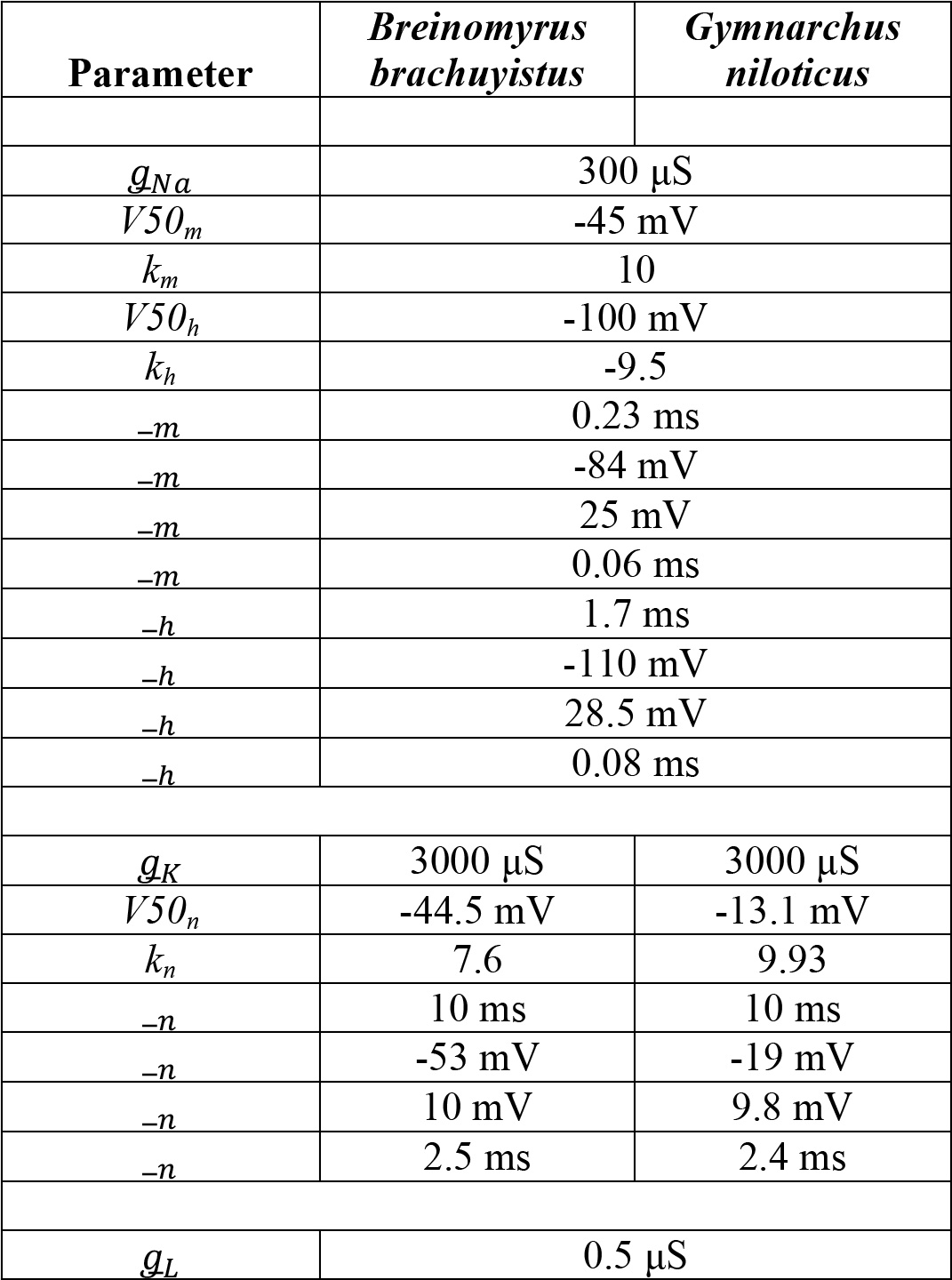
Parameter values for initial electrocyte models.

**Table S4.**
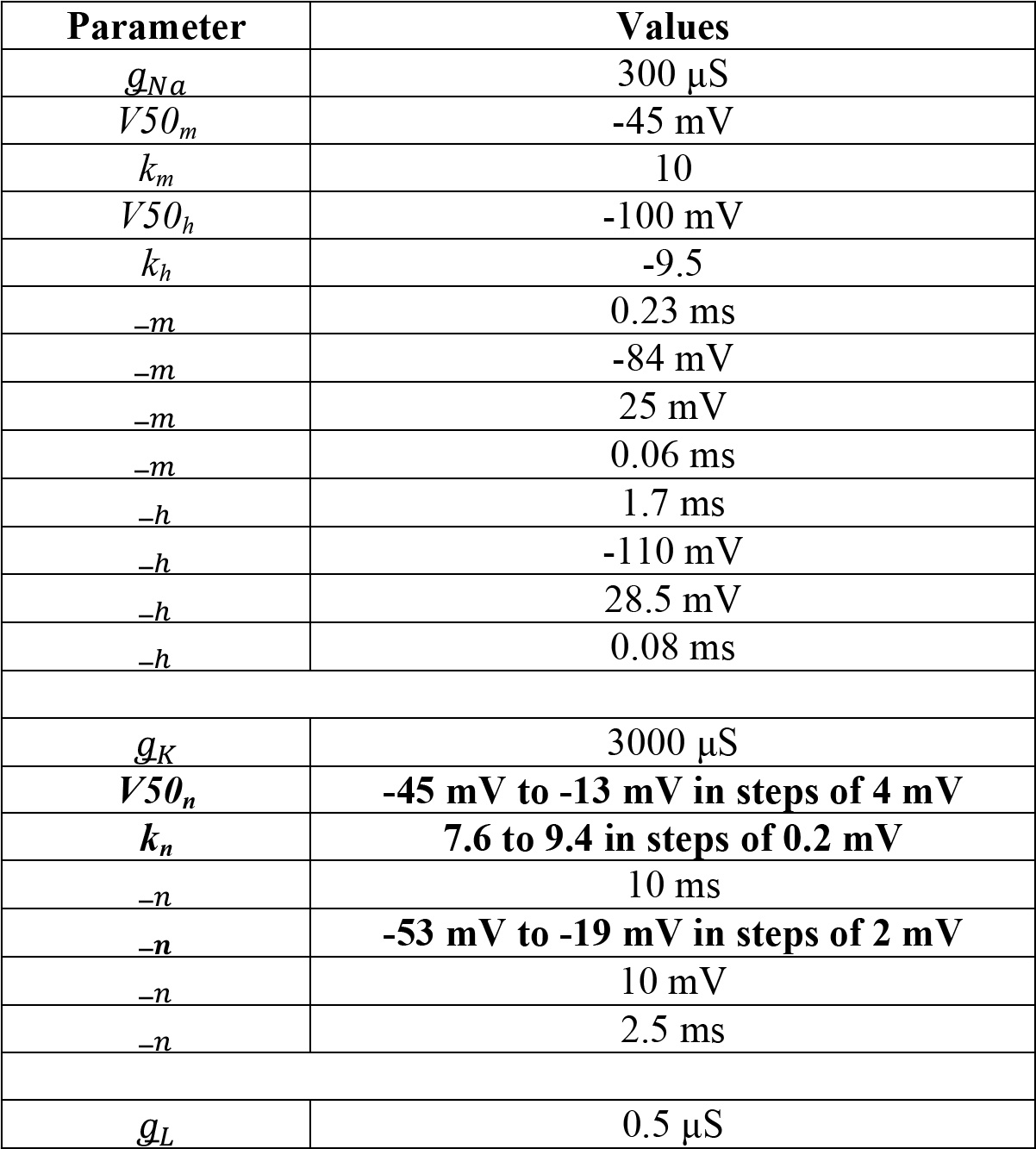
Parameter values for intermediate electrocyte models.

**Table S5:**
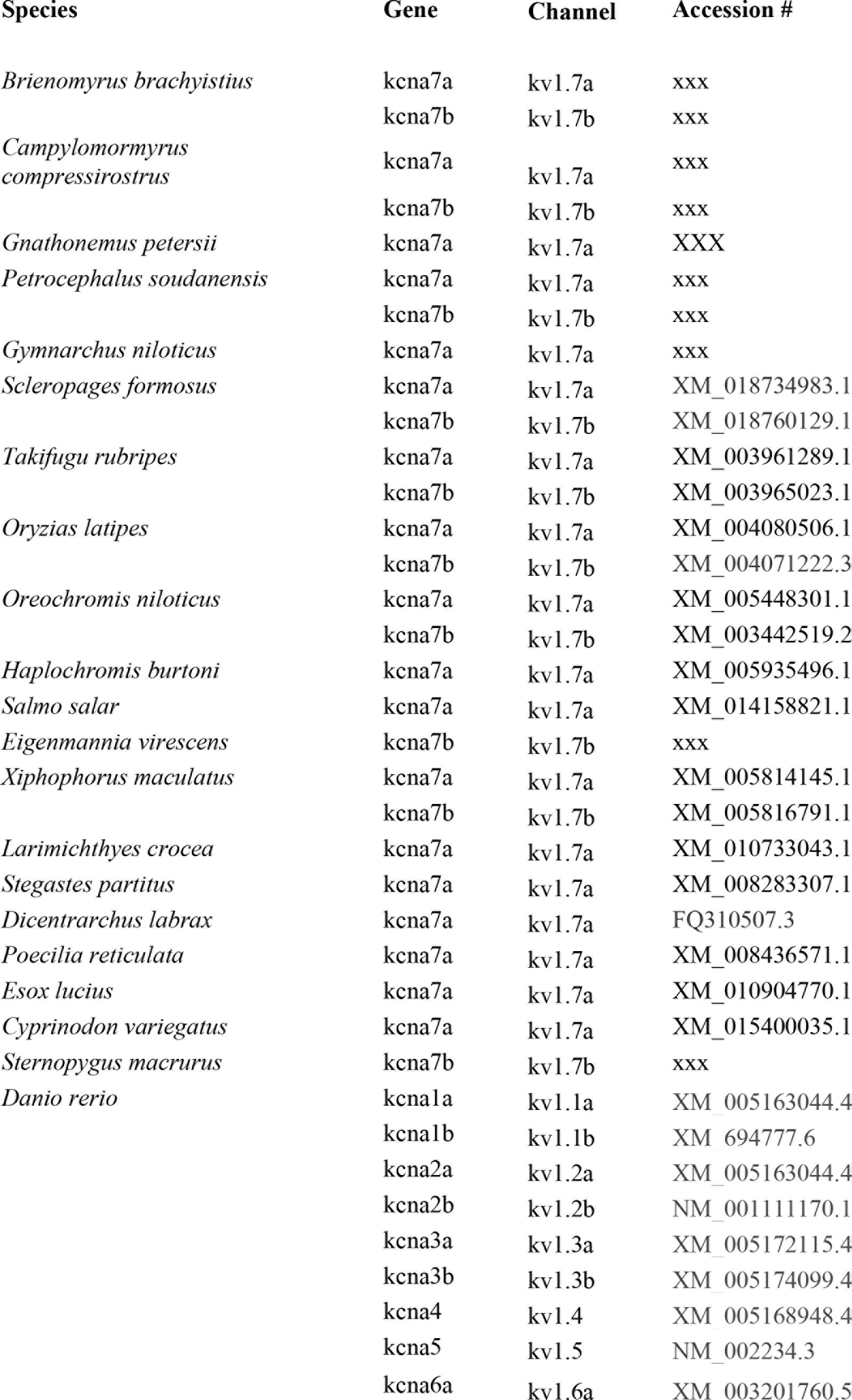
Sequences and accession numbers of genes for phylogenenetic trees in figs.1 and.

**Table S6:**
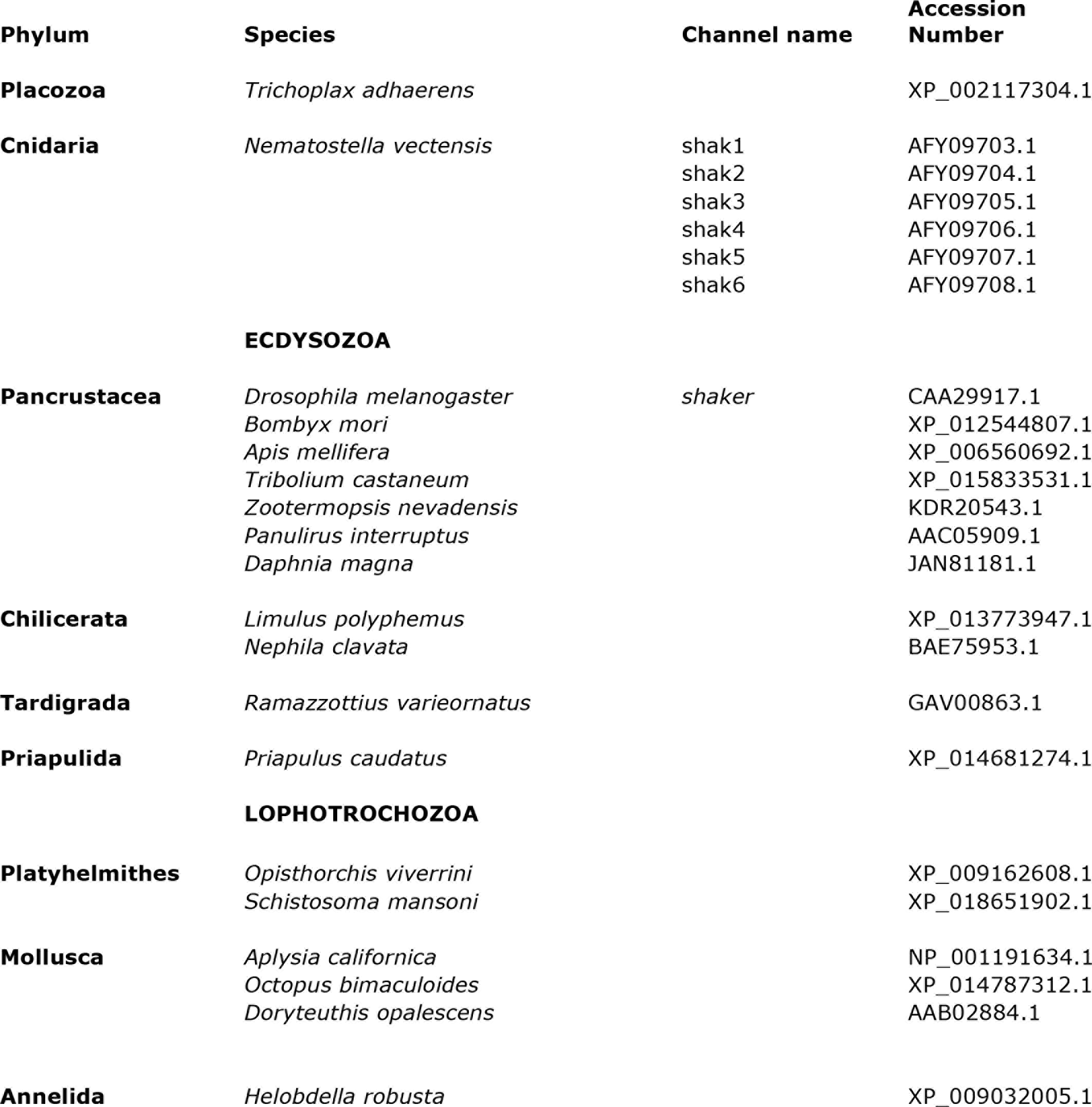
Species and accession numbers of genes used for analysis of S3-S4 linkers in Fig. S6.

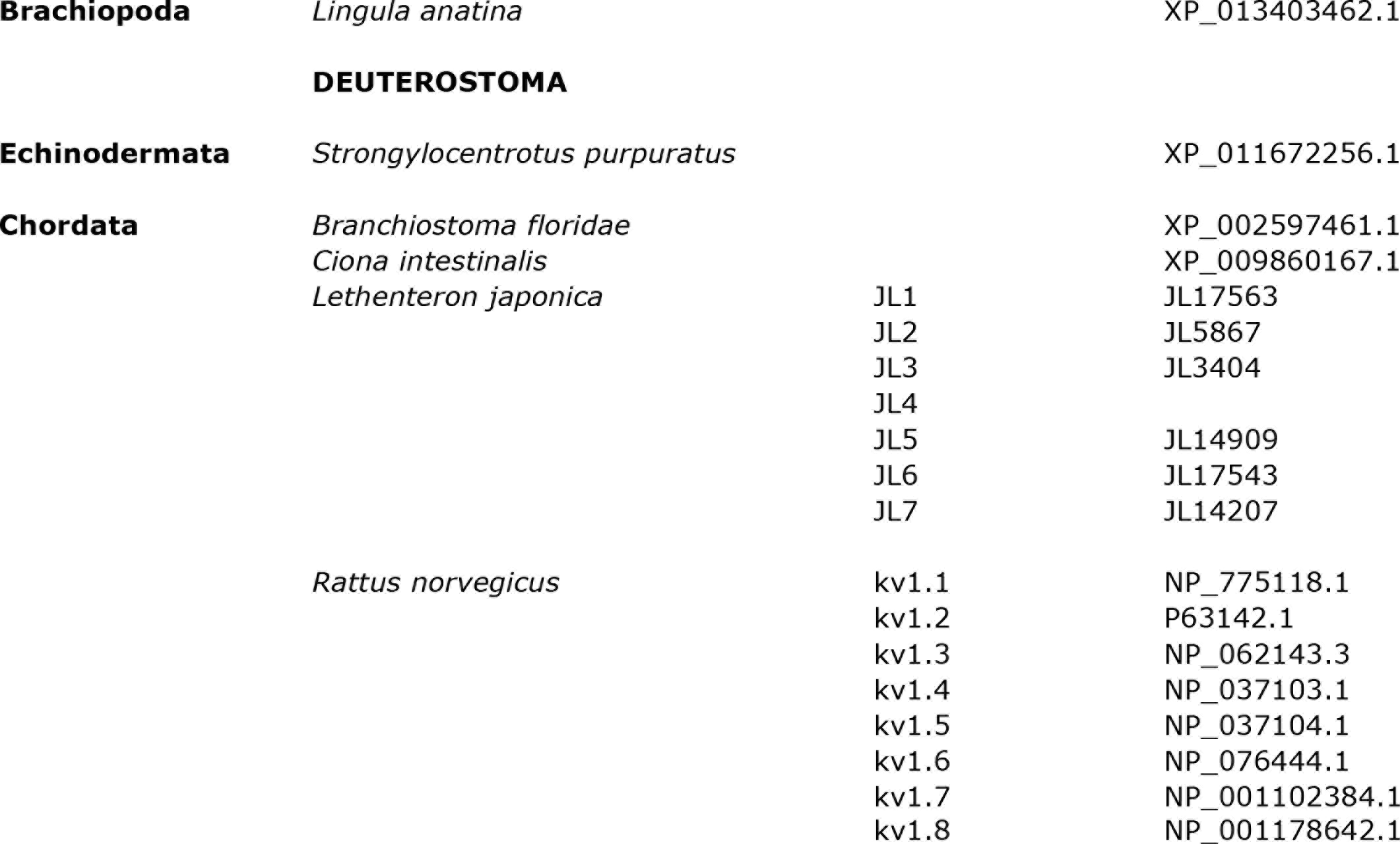

